# Ecological networks balance connectivity with flexibility to perturbations

**DOI:** 10.64898/2026.06.12.731932

**Authors:** Àlex Giménez-Romero, Daniel Oro, Jordi Bascompte, Meritxell Genovart

**Affiliations:** Department of Ecology and Complexity, Centre d’Estudis Avançats de Blanes (CEAB-CSIC), Blanes, Girona, Spain; Department of Evolutionary Biology and Environmental Studies, University of Zurich, Zurich, Switzerland

## Abstract

Food webs become progressively sparser as species richness increases, yet the number of interactions grows faster than linearly, placing communities between the limiting expectations of constant degree and constant connectance. Similar scaling has also been documented in other interaction networks, but whether they share a common architectural regime, and what general principle could generate it, remain unresolved. Here, we show that 1,533 ecological networks spanning different interaction types broadly converge on an intermediate link–species scaling regime, with an overall exponent of 1.43. We interpret this pattern through a thermodynamic trade-off in which increasing network connectivity enhances signal propagation while reducing the diversity of independent modes contributing to the network’s response to perturbations. Optimizing this trade-off yields a scaling exponent of 1.45, closely matching the empirical estimate, while randomly adding or removing interactions generally reduces thermodynamic efficiency in empirical networks. These results support a propagation–response-diversity trade-off as a plausible systems-level principle underlying how ecological network complexity grows with diversity.

## Introduction

Ecological communities are not simply collections of species, but networks of interactions whose architecture determines how ecological complexity grows with diversity. For more than four decades, ecologists have therefore asked how the number of interactions, *L*, increases with species richness, *S*. The answer to this question is central to ecology because it determines how increases in species richness, such as those occurring during ecological succession after perturbation, translate into changes in the structure and dynamics of ecological communities. Early analyses of food webs suggested a surprisingly simple organization: the number of trophic links per species appeared approximately invariant across communities [1, 2], giving rise to the link–species scaling law *L* ∝ *S*. This apparent scale invariance became one of a broader set of proposed regularities in food-web architecture [3–5]. As increasingly resolved webs became available, however, these regularities proved controversial. Analyses of detailed empirical networks showed progressively more links per species as richness increased [6–9], while others suggested instead that connectance—the fraction of possible interactions that are realized—remained approximately invariant [10, 11], implying the much faster scaling *L* ∝ *S*^2^. Subsequent work showed that inferred relationships also depend on sample size, taxonomic resolution, aggregation, and sampling effort [12–15], making the relationship between interaction richness and species richness one of the most persistent conceptual conundrums in food-web ecology [16].

Improved and increasingly quantitative datasets have since converged on a more nuanced picture. Empirical food webs generally occupy neither of the two classical limits: links per species tend to increase with richness, while connectance decreases, such that the number of interactions grows faster than linearly but slower than quadratically [17–20]. The resulting intermediate relationship, *L* ∝ *S^γ^* with 1 *< γ <* 2, has therefore emerged repeatedly despite substantial differences among food webs. Yet the scaling exponent is not itself an explanation. Several mechanisms can constrain how interactions accumulate with diversity. Optimal-foraging and diet-breadth constraints can reproduce non-trivial connectance scaling in trophic networks [21]; spatial processes couple species accumulation to the accumulation of interactions across area [22–24]; and physical or physiological constraints restrict the set of interactions that can be realized [25]. Classical complexity–stability theory provides an additional reason why increasing community size may favor relative sparsity [26, 27], while more recent theory has connected link–species relationships themselves to robustness and local stability [28]. These perspectives identify mechanisms that generate network structure and consequences of that structure for stability and robustness. A complementary question is why evolutionary and ecological processes should repeatedly favour architectures with intermediate link–species scaling.

The scope of this problem also extends beyond food webs. Declining connectance with increasing richness was reported early for plant–pollinator and seed-dispersal networks [29, 30], and interaction richness has subsequently been shown to increase non-trivially with diversity or spatial extent in pollination, host–parasite, and other ecological networks [23, 24, 31, 32]. Ecological-network research has consequently moved increasingly towards identifying principles that transcend particular forms of interaction and allow network structure to be understood or predicted across ecological contexts [33, 34]. Yet existing evidence remains fragmented among mutualistic and antagonistic systems, or derives from networks assembled along particular spatial gradients with the estimated scaling relationships varying among datasets [24]. It therefore remains unclear whether fundamentally different ecological interaction types converge on a common intermediate link–species regime when examined within the same comparative framework. Establishing such a regularity would be consequential: despite profound differences in the biological mechanisms determining who interacts with whom, a shared scaling regime would point towards constraints on network organization that operate above the level of particular interaction types.

A candidate explanation emerges from considering what connectivity does at the level of the whole network. Ecological interactions provide pathways through which environmental perturbations, energy, matter and information can propagate across a community [35–37]. Increasing connectivity can therefore enhance integration among network components, but stronger coupling also makes their responses progressively less independent, reducing the diversity of dynamical modes available to the system. Such response diversity can be important for ecological functioning and stability because heterogeneous responses limit the extent to which perturbations propagate through ecological communities, thereby reducing the risk of collapse [38]. Related ideas have also emerged from studies of animal social networks, where environmental perturbations can modify network cohesion and alter how responses are distributed across individuals [39]. A recent framework from the statistical physics of information dynamics formalizes this tension through diffusion on networks [40–42]. In this framework, the Laplacian spectrum determines both how signals propagate through the network and the diversity of independent modes contributing to its response. Their balance can be quantified through a measure of thermodynamic efficiency that combines gains in propagation with losses in the diversity of responses to perturbations, and whose optimization generates sparse architectures [42]. Unlike mechanisms tied to a particular ecological interaction, this principle depends primarily on the collective consequences of network coupling, providing a potential systems-level explanation for both the recurrence of intermediate connectivity and its apparent generality across ecological networks.

Here, we explore both the generality and possible origin of the intermediate link–species scaling using 1,533 empirical ecological networks spanning different interaction types. We first ask whether fundamentally different interaction types share a common architectural regime, and show that interaction richness increases superlinearly with species richness, globally as *L* ∼ *S*^1.43^, while connectance declines and the number of interactions per species increases. We then ask whether this regime is consistent with a trade-off between perturbation propagation and response diversity. Randomly adding or removing interactions generally reduces thermodynamic efficiency, while the stationary condition of an analytical approximation yields a link-scaling exponent close to that observed empirically. Finally, degree-preserving null models reveal how higher-order network organization modulates this balance among interaction types. Together, these analyses show that a propagation–response-diversity trade-off can provide a common organizing principle for how ecological network complexity grows with diversity and sheds light on how ecological communities might respond to environmental perturbations.

### A common intermediate scaling regime across ecological interaction networks

To determine whether link–species scaling follows a common pattern across contrasting ecological interactions, we assembled 1,533 empirical networks from the Web of Life and Mangal databases, spanning different interaction types across terrestrial, freshwater, and marine environments (Fig. 1a). Across the full dataset, the number of interactions increased superlinearly with species richness,

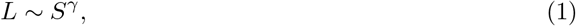

with *γ* = 1.43 (95% CI: [1.41, 1.45], *R*^2^ = 0.93, *p <* 0.001; Fig. 1c).

**Figure 1:**
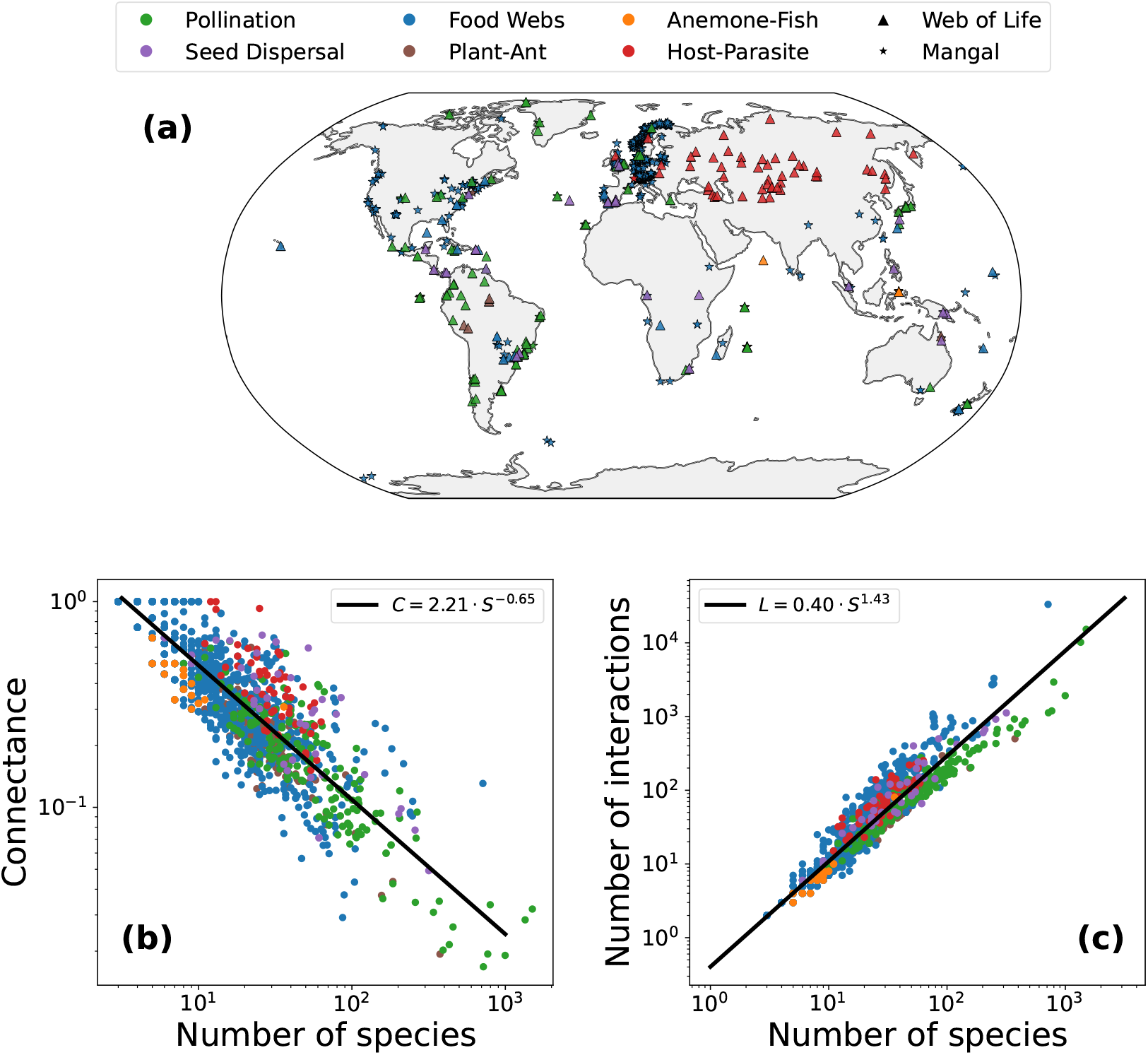
Ecological interaction networks occupy an intermediate link–species scaling regime. **(a)** Geographic distribution of the 1,533 ecological interaction networks compiled from the *Web of Life* and *Mangal* databases. Colors indicate interaction type and marker shapes denote the source database. The dataset spans terrestrial, freshwater, and marine ecosystems across multiple continents. **(b)** Connectance declines systematically with species richness, *C* ∼ *S*^−0.65^ (95% CI: [−0.67,−0.64], *R*^2^ = 0.78, *p <* 0.001). Points are colored by interaction type and the dashed line denotes the global power-law fit. **(c)** The total number of interactions increases superlinearly with species richness, *L* ∼ *S^γ^*, with *γ* = 1.43 (95% CI: [1.41, 1.45], *R*^2^ = 0.93, *p <* 0.001). The observed exponent lies between the limiting expectations of constant degree (*L* ∼ *S*) and constant connectance (*L* ∼ *S*^2^), defining an intermediate scaling regime.

Ecological communities therefore accumulate interactions faster than expected if species maintained a constant number of partners (*L* ∼ *S*), but substantially more slowly than the quadratic growth expected under constant connectance (*L* ∼ *S*^2^).

This intermediate regime is expressed simultaneously through decreasing interaction density and increasing interactions per species. Connectance declined systematically with richness as *C* ∼ *S*^−0.65^ (95% CI: [−0.67,−0.64], *R*^2^ = 0.78, *p <* 0.001; Fig. 1b), whereas mean degree increased as ⟨*k*⟩ ∼ *S*^0.43^ (*R*^2^ = 0.54, *p <* 0.001; Supplementary Fig. 1c). Thus, species in richer communities interact, on average, with progressively more partners, even though those interactions represent a decreasing fraction of all possible links. The observed architecture consequently differs from both classical limiting cases: neither a fixed interaction capacity per species nor constant connectance describes how ecological complexity accumulates with diversity [11, 43, 44].

Importantly, this intermediate scaling was not generated by pooling structurally distinct ecological networks. When each interaction class was analyzed separately, interaction richness increased superlinearly with species richness in every case, with exponents ranging from *γ* = 1.22 for plant– pollinator networks to *γ* = 1.63 for food webs (Supplementary Fig. 3). All confidence intervals lay between the limiting expectations *γ* = 1 and *γ* = 2, showing that networks with different interaction types independently occupy the same intermediate link–species regime, although the rate at which interactions accumulate varies among them. This distinction also persisted when networks were grouped by their structural representation: bipartite networks followed *L* ∼ *S*^1.34^ (*R*^2^ = 0.96, *p <* 0.001; Supplementary Fig. 2a), whereas unipartite networks scaled more steeply, *L* ∼ *S*^1.47^ (*R*^2^ = 0.85, *p <* 0.001; Supplementary Fig. 2b). Thus, the intermediate regime is shared across ecological networks, while its quantitative exponent depends on interaction type.

The same pattern can be viewed through its complementary consequences for interaction density and interactions per species. Connectance declined significantly with richness within every interaction class considered (Supplementary Fig. 4), while mean degree increased with richness across all interaction types (Supplementary Fig. 8). Notably, the diversity–connectance relationship was more conserved across interaction classes than the corresponding link–species exponents and remained similar when bipartite and unipartite networks were analysed separately (Supplementary Fig. 5). This greater consistency is compatible with connectance being normalized by the relevant space of possible interactions (*S*(*S* − 1)*/*2 for unipartite networks and *S*_1_*S*_2_ for bipartite networks) thereby accounting for fundamental differences in how potential interactions are defined across network types. These common trends emerge despite marked differences among interaction classes in their typical size, connectance, mean degree, degree heterogeneity, and modularity (Supplementary Table 3). Ecologically distinct networks therefore share a common tendency towards increasing relative sparsity with diversity, while differing in the rate at which absolute interaction richness accumulates.

Two established perspectives provide a useful reference for this intermediate regime, but neither explains it quantitatively. Classical complexity–stability theory predicts that sufficiently large communities may require decreasing connectance, and several random-matrix formulations recover the asymptotic scaling *C* ∝ *S*^−1^. However, these results depend on assumptions about interaction strengths, self-regulation, and network structure, and are therefore best regarded here as theoretical benchmarks rather than direct predictions for the empirical networks (Supplementary Section 1). At the opposite structural limit, the constant-degree hypothesis predicts *L* ∝ *S*, because each species retains an approximately fixed number of interaction partners as richness increases [43, 44]. Our data depart from both expectations: connectance declines more slowly than *S*^−1^, while mean degree increases systematically with richness and degree distributions remain strongly heterogeneous (Supplementary Fig. 7).

Taken together, these results reveal a specific scaling regime in ecological network architecture. Connectance decreases with richness, *C* ∼ *S*^−0.65^, yet average degree increases, ⟨*k*⟩ ∼ *S*^0.43^, implying that the total number of interactions grows superlinearly with community size, *L* ∼ *S*^1.43^. Ecological communities therefore lie between two limiting cases: the linear scaling expected under the constant-degree hypothesis (*L* ∼ *S*) and the quadratic scaling of a fully connected network (*L* ∼ *S*^2^). Neither classical stability theory alone nor simple topological null arguments explain why ecological systems consistently occupy this intermediate regime. Instead, this recurring pattern suggests that ecological communities may be organized within a broader structural envelope shaped by multiple constraints.

### A thermodynamic basis for intermediate link–species scaling

We next asked whether the intermediate link–species scaling observed across ecological networks could arise from a general constraint associated with network connectivity. Ecological interactions provide pathways through which both beneficial and detrimental effects can spread across a community. On the one hand, they enable species to access energy and other resources and to acquire information, supporting ecological functioning and integration [35, 37]. On the other hand, the same interactions can transmit disturbances, such as disease or demographic changes, potentially affecting species far beyond those initially impacted [36]. Increasing connectivity can therefore facilitate transport across the community, but also expose more species to disturbances. Ecological networks may thus face a trade-off between maintaining efficient transport and preserving the community’s capacity to respond to perturbations in multiple, independent ways.

A recent thermodynamic framework for diffusion on complex networks provides a way to quantify this trade-off [42]. The central idea is to consider how a quantity initially localized at one species spreads through the network. As in classical diffusion, this quantity progressively redistributes across the system, but here propagation occurs along network links rather than through physical space. A signal initially localized at one species can spread through the network, first reaching its direct interaction partners and then propagating to other species along successive links. Diffusion provides a minimal description of this propagation process, without specifying the particular ecological mechanisms involved. However, understanding how a signal spreads from a single species is not sufficient to characterize the collective response of the community. Signals originating at different species may propagate through different pathways, affecting different parts of the network and producing distinct or overlapping response patterns. The framework therefore considers all species as potential origins of a signal and combines their individual responses into a statistical description of the entire network, known as the network density matrix (see Methods). This representation captures both how signals are distributed across species and how their responses are correlated, allowing us to characterize the network’s collective propagation properties across different scales.

Within this statistical physics framework, two complementary quantities characterize the effects of network connectivity. The network free energy measures the capacity of the network to transport signals among its components [45], with higher values corresponding to more efficient propagation and less local trapping. By contrast, the von Neumann entropy [46] measures the diversity of diffusion modes contributing to the network’s response [40]. These modes represent different collective patterns through which signals can spread and relax across the community. High entropy indicates that many distinct modes contribute to the response, whereas low entropy indicates that the response is dominated by fewer collective modes. Relative to a reference network, increasing connectivity enhances signal propagation while reducing response diversity [42] (see Methods). These opposing effects can be quantified as a gain in free energy, *W_τ_*, and an entropy-associated cost, *Q_τ_*, which mathematically resemble work and heat in thermodynamics, respectively. Their relative balance defines the thermodynamic efficiency [42],

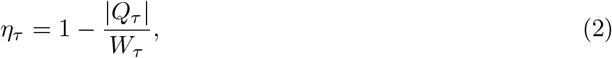

where *τ* represents the propagation scale (see Methods). High thermodynamic efficiency thus reflects a favourable balance between transport capacity and response diversity, rather than the maximization of either property independently.

This trade-off is illustrated in Fig. 2. Across random networks of fixed size, increasing connectance monotonically enhances perturbation propagation but progressively reduces the diversity of response modes (Fig. 2a). Their combination produces a maximum in thermodynamic efficiency at intermediate connectivity. The same pattern is visible in individual network realizations. In sparsely connected networks, an initially localized perturbation remains unevenly distributed across nodes over a broad range of propagation scales, preserving high response diversity but limiting system-wide propagation (Fig. 2b,e,h). Increasing connectivity allows the perturbation to spread more broadly while responses remain differentiated, generating the highest efficiency at intermediate connectivity (Fig. 2c,f,i). At still higher connectivity, responses rapidly converge towards the homogeneous state: propagation is maximized, but response diversity is largely lost and efficiency consequently declines (Fig. 2d,g,j). Thus, the efficiency maximum arises from the opposing dependence of propagation and response diversity on network connectivity.

**Figure 2:**
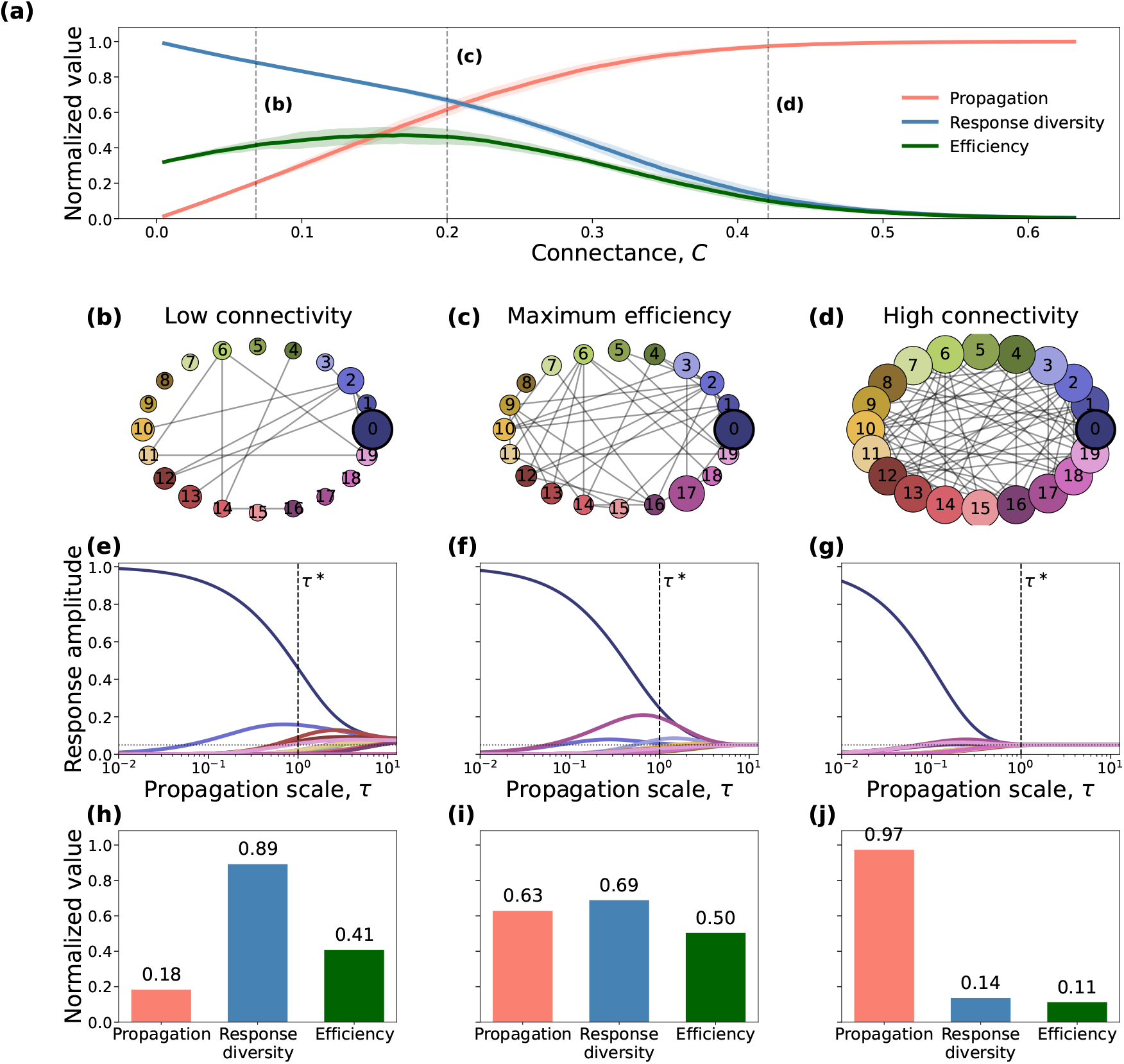
Thermodynamic trade-off between perturbation propagation and response diversity. **(a)** Normalized propagation, response diversity, and thermodynamic efficiency (the balance between the gain in propagation and the associated loss of response diversity) as a function of network connectance for random networks with *N* = 20 nodes. Lines show the mean across random realizations and shaded regions indicate ±1 s.d. Propagation increases with connectance, whereas response diversity decreases, producing an intermediate-connectance maximum in thermodynamic efficiency. Vertical dashed lines indicate the connectances of the three representative networks shown in **b–d**. **(b–d)** Representative low-connectivity, maximum-efficiency, and high-connectivity networks, respectively. Node colors identify individual nodes across panels, and node size is proportional to the response amplitude at the reference propagation scale *τ* ^∗^ normalized to the maximum response value. **(e–g)** Diffusive response of each node to a perturbation initially localized at node 0 as a function of propagation scale *τ*. Colors correspond to the nodes in **b–d**. The vertical dashed line denotes *τ* ^∗^ = 1, at which the thermodynamic quantities are evaluated, and the horizontal dotted line indicates the homogeneous response 1*/N*. Increasing connectivity enables the perturbation to propagate more rapidly through the network but progressively reduces differentiation among node responses. **(h–j)** Normalized propagation, response diversity, and thermodynamic efficiency evaluated at *τ* ^∗^.

This framework provides a quantitative connection to the link–species scaling observed empirically. Suppose that interaction richness scales with species richness as

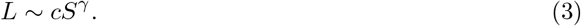

Within the mean-field approximation of the thermodynamic efficiency, the stationary condition with respect to connectivity yields an effective exponent

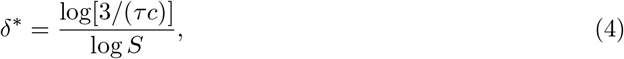

with the corresponding link–species exponent

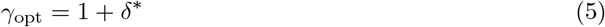

(see Methods). Evaluating this condition for each empirical network at its characteristic diffusion scale, *τ*_diff_ = 1*/λ*_2_, gives ⟨*γ*_оpt_⟩ = 1.45 (Fig. 3), remarkably close to the empirical estimate *γ* = 1.43 (95% CI: [1.41, 1.45]; Fig. 1). This correspondence was also recovered when bipartite and unipartite networks were analyzed separately. The efficiency approximation yielded ⟨*γ*_оpt_⟩ = 1.37 and 1.43, respectively, close to the corresponding empirical exponents *γ* = 1.34 and 1.47 (Supplementary Fig. 15).

**Figure 3:**
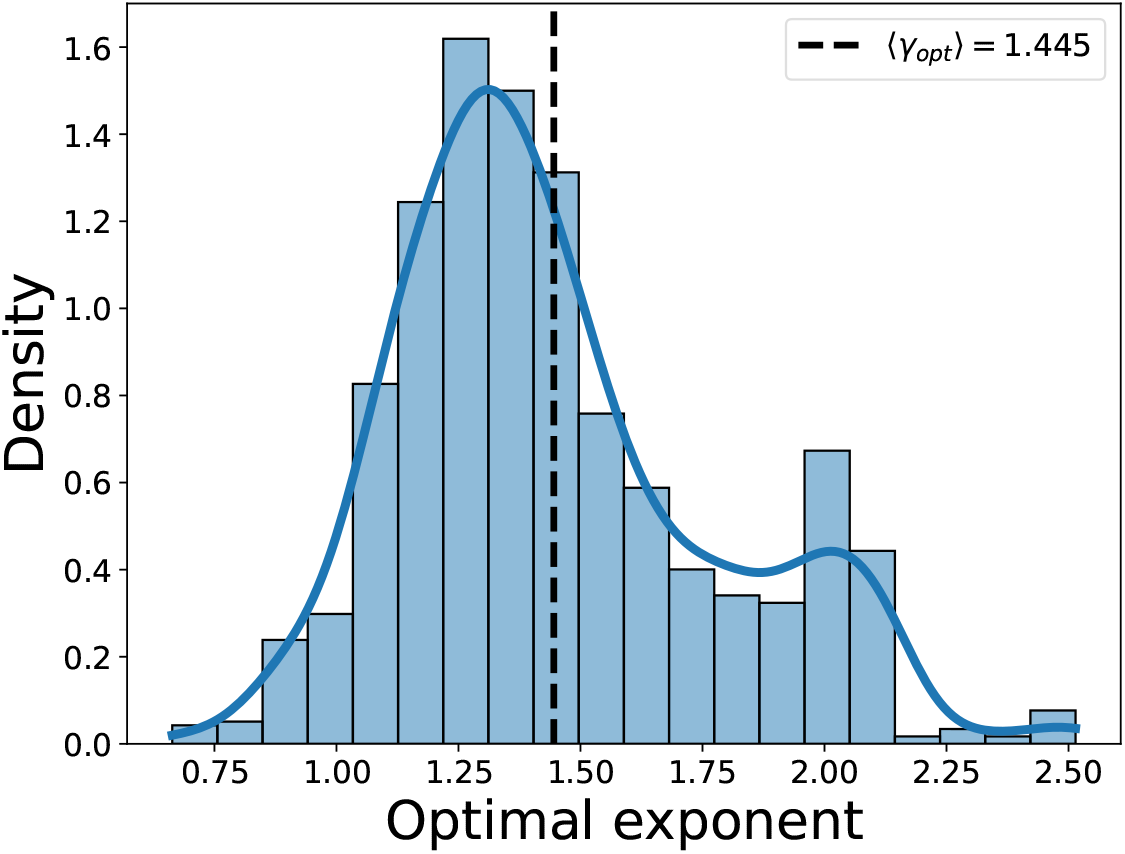
Distribution of optimal scaling exponents from the efficiency approximation. Distribution of the optimal scaling exponent *γ*_оpt_ = *δ*^∗^ + 1 obtained by evaluating the stationary condition of the thermodynamic efficiency approximation for each empirical network at the characteristic diffusion scale *τ*_diff_ = 1*/λ*_2_. The average value of ⟨*γ_opt_*⟩ = 1.445 is in striking agreement with the empirically fitted scaling exponent *γ* = 1.43 [1.41, 1.45] obtained from the fit *L* ∼ *S^γ^*. This agreement indicates that the observed super-linear scaling of interactions with species richness is consistent with the balance between propagation and response diversity captured by the thermodynamic efficiency framework.

Given the mean-field approximation and the finite size and structural heterogeneity of the empirical networks, this agreement should not be interpreted as a parameter-free derivation of the observed exponent. It nevertheless provides a direct quantitative link between the propagation– response-diversity trade-off and the intermediate scaling regime observed across ecological interaction networks: the stationary point of the thermodynamic efficiency occurs at essentially the same rate of interaction accumulation as that measured empirically.

### Thermodynamic efficiency in ecological networks

The close correspondence between the efficiency-based stationary condition and the empirical link–species exponent suggests that the propagation–response-diversity trade-off may help constrain how connectivity scales with diversity. Yet this scaling-level correspondence says little about the architecture of individual communities. We therefore first examined how thermodynamic efficiency is expressed across empirical ecological networks. Despite their substantial structural differences, all interaction types exhibited the same qualitative efficiency profile (Fig. 4a, Supplementary Fig. 10): efficiency was high at small propagation scales, when perturbations remained localized, and declined as increasing *τ* progressively integrated the network and reduced the diversity of independent response modes. The characteristic scale of this decline nevertheless differed among interaction types, indicating systematic variation in structural coupling.

**Figure 4:**
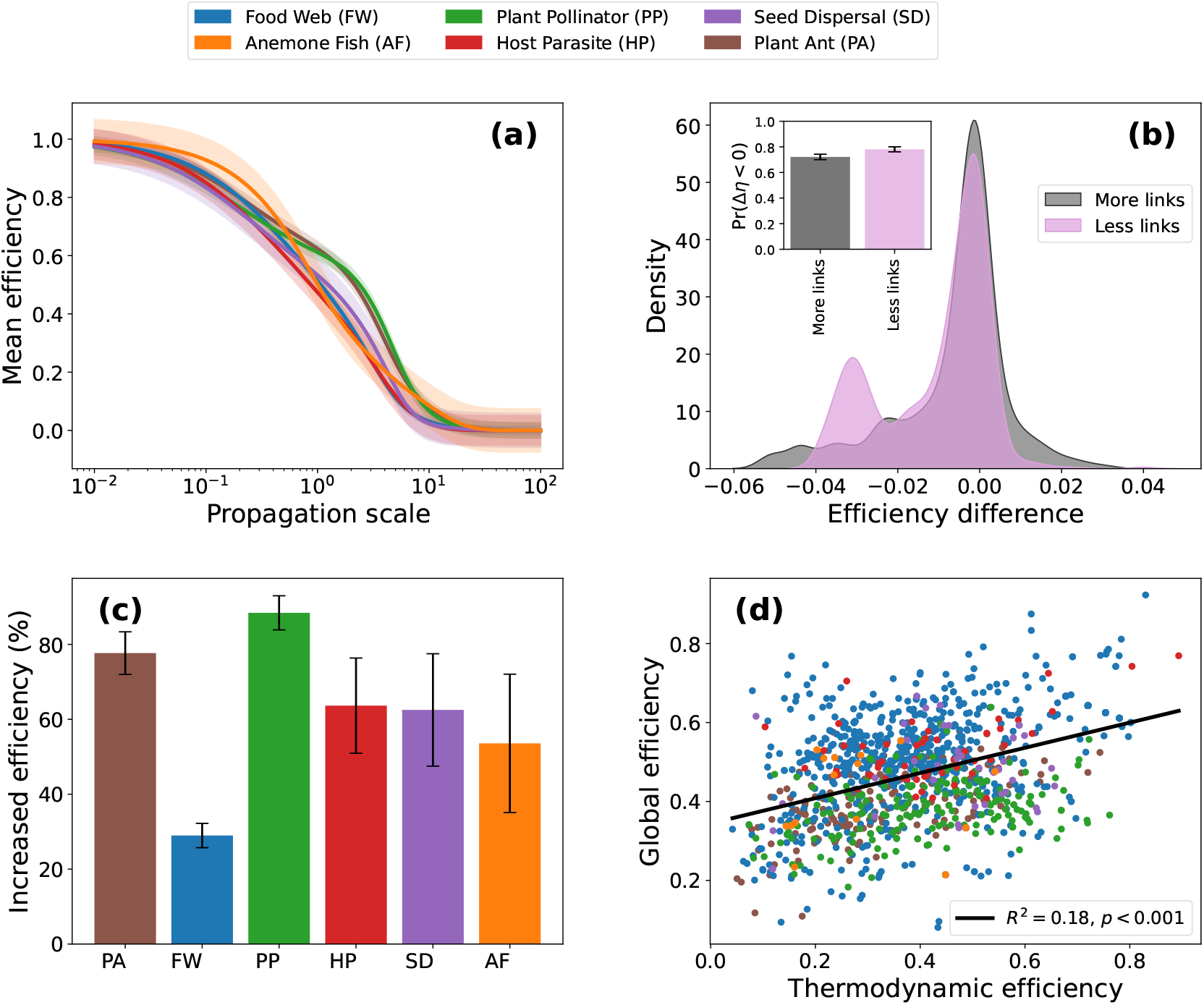
Thermodynamic efficiency of ecological interaction networks. **(a)** Mean efficiency profiles ⟨*η*(*τ*)⟩ for each interaction type, highlighting differences in how efficiency declines with propagation scale. Individual profiles are shown in Supplementary Fig. 10, where differences among individual networks are highlighted. **(b)** Distribution of differences in thermodynamic efficiency between empirical networks and perturbed networks constructed by increasing (black) or decreasing (pink) the number of interactions by 1%. For each network, differences were computed across the propagation range *τ* ∈ [10^−2^, 10^2^] and averaged over *τ*. The distribution indicates that empirical networks lie near an optimal balance between propagation and response diversity, such that small increases or decreases in interaction density tend to reduce thermodynamic efficiency. (**c**) Fraction of networks within each interaction type exhibiting substantial positive deviations in thermodynamic efficiency relative to the configuration-model (CM) null ensemble, defined as Δ*η*_CM_ *>* 0.1. Error bars denote binomial 95% confidence intervals. Such deviations are particularly frequent in mutualistic networks, especially plant–pollinator and plant–ant communities, whereas food webs exhibit them much less frequently. (**d**) Relationship between thermodynamic efficiency and standard global efficiency (mean inverse shortest-path length). The positive association (*R*^2^ = 0.18, *p <* 0.001) indicates that networks with increased thermodynamic efficiency also tend to exhibit enhanced topological efficiency.

Whether these profiles reflect distinctive structural organization, however, cannot be established from the empirical networks alone. We therefore compared them with null ensembles imposing different constraints on network architecture. Erdös–Rényi networks preserving network size and expected interaction density exhibited systematically lower efficiency at intermediate propagation scales than their empirical counterparts (Supplementary Fig. 11, Supplementary Fig. 12a), showing that interaction density alone does not account for the observed profiles. Configuration-model (CM) networks accounting for empirical degree heterogeneity more closely reproduced the diffusion behavior (Supplementary Fig. 11), but substantial positive efficiency deviations remained across interaction types (Supplementary Fig. 12b), particularly in plant–pollinator networks. These results indicate that species-level degree heterogeneity explains part of the empirical efficiency profiles, but does not fully account for the observed thermodynamic efficiency.

The magnitude of these deviations from CM expectations was not uniformly distributed across interaction types. In particular, plant–pollinator and plant–ant networks frequently exhibited substantial positive efficiency deviations (Δ*η*_CM_ *>* 0.1), whereas such large deviations were considerably less common in food webs (Fig. 4c). However, when the empirical degree sequence of bipartite networks was preserved exactly using the fixed-marginal Curveball null model [47], efficiency deviations were substantially reduced and generally concentrated around zero (Supplementary Fig. 13). This result should nevertheless be interpreted cautiously, as the structural constraints imposed by null models can strongly influence the inferred deviations from empirical network organization [48–50]. In particular, preserving the degree sequence exactly may severely restrict the space of admissible network configurations, potentially leaving randomized networks structurally similar to the empirical ones.

These comparisons show that empirical network structure carries a non-trivial efficiency signature, but they do not establish whether the observed architectures themselves lie near a favorable point of the efficiency landscape. If the propagation–response-diversity trade-off helps shape connectivity at the level of individual networks, departures from the observed interaction density should tend to reduce efficiency. We therefore randomly added or removed interactions from each empirical network. Changing the number of interactions by only 1% produced predominantly negative differences in efficiency in both directions (Fig. 4b). This pattern persisted across perturbations of 1–10% of empirical interaction richness (Supplementary Fig. 14), for which both link addition and removal consistently reduced efficiency. Empirical networks are therefore locally consistent with a maximum of the thermodynamic-efficiency metric: additional interactions improve propagation but accelerate the loss of independent response modes, whereas removing interactions preserves response diversity at the expense of propagation. Their observed connectivity lies between these opposing tendencies, where the balance between propagation and response diversity is locally most favorable.

To further characterize the relationship between network architecture and efficiency, we compared thermodynamic efficiency with standard global efficiency (mean inverse shortest-path length). Networks with higher thermodynamic efficiency also tended to exhibit higher global efficiency (Fig. 4d; *R*^2^ = 0.18, *p <* 0.001), indicating a positive, although relatively weak, association between the propagation–response-diversity trade-off and the accessibility of species through shorter interaction pathways. Together, these results suggest that while the overall sparsity scaling is shared across interaction types, species-level connectivity and higher-order network organization contribute differently to thermodynamic efficiency. Thus, within the common architectural constraint linking richness and connectance, interaction type may modulate how effectively networks balance coupling and response diversity.

## Discussion

Ecological networks gain interactions as they gain species, but at a remarkably constrained rate. Across 1,533 networks spanning different interaction types, interaction richness scales as *L* ∼ *S*^1.43^: the average number of partners per species increases with diversity, while the fraction of all possible interactions declines. This places ecological communities between the two classical limits of constant degree (*L* ∝ *S*) and constant connectance (*L* ∝ *S*^2^) [1, 11]. Importantly, this intermediate regime recurs across ecologically distinct interaction types, despite differences in their scaling exponents and in the ecological and evolutionary processes that generate individual interactions. Such convergence suggests that interaction-specific mechanisms alone are unlikely to determine how connectivity scales with diversity. Similar constraints on network architecture have been identified in other complex systems. For example, brain networks balance efficient information transfer against wiring costs [51], while transportation networks face trade-offs between connectivity and infrastructure overheads [52]. More generally, the balance between information propagation and response diversity has been proposed as an explanation for the sparsity of real-world networks across different domains [42]. Together, these observations suggest that, while ecological mechanisms determine the interactions between species, broader system-level constraints may govern how connectivity scales with diversity across ecological networks.

The propagation–response-diversity trade-off suggests what one such constraint could be. Every additional interaction creates a new route through which effects can move across a community, while also coupling the responses of previously more independent components. Increasing connectivity therefore improves network integration at the cost of progressively reducing the diversity of independent response modes [42]. Our empirical networks bear both signatures of the resulting balance. The stationary condition of the analytical approximation gives ⟨*γ*_оpt_⟩ = 1.45, close to the observed *γ* = 1.43, and randomly adding or removing interactions from individual networks generally lowers thermodynamic efficiency. The first result links the trade-off to how connectivity changes across communities of different richness; the second shows that the observed interaction density of individual communities tends to lie in a locally favorable region of the efficiency landscape. The quantitative agreement is not a parameter-free derivation of the scaling law, nor do local perturbations establish a global optimum. Together, however, they suggest that the intermediate scaling of ecological networks is consistent with a balance between the benefits and costs of increasing network coupling. This balance may ultimately have an evolutionary basis, provided that ecological and evolutionary processes have repeatedly favored network architectures that maintain both integration and flexibility.

There is an ecological basis for both sides of this balance. Connectivity is both opportunity and exposure. Species interactions convey energy, matter and information, allowing local changes to influence distant parts of a community and thereby increasing network-wide integration [35–37]. At the same time, differentiated responses are themselves ecologically valuable. Species that respond differently to environmental change generate asynchronous population dynamics, buffering aggregate community properties through time. Recent work in complex food webs identifies response diversity as a major driver of temporal stability through precisely this effect on population asynchrony [38]. The response diversity quantified here is spectral rather than species-specific: it describes the structural modes through which perturbations can propagate, rather than variation in environmental response traits. Yet the underlying systems property is closely related. Stronger coupling integrates the community while making its components respond less independently. Intermediate connectivity likely preserves substantial propagation without collapsing the response of the system onto a small number of collective modes.

This does not imply that ecological communities evolve to maximize thermodynamic efficiency itself. Natural selection acts on heritable variation affecting survival and reproduction [53], but the fitness consequences of that variation depend partly on interactions with other species. The ecological and evolutionary trajectories of interacting species are therefore interdependent. We propose that selection on organismal traits, together with the population dynamics and ecological dependencies linking species, can contribute to the emergence of community architecture without requiring direct selection on its collective properties. Thus, high thermodynamic efficiency may be an emergent outcome of these coupled processes. Interactions carry benefits and costs for the species involved, and recent eco-evolutionary models show that trade-offs in acquiring interactions can generate distinct connectivity regimes during community assembly [54]. Assembly also filters interaction structures according to whether they can support persistent communities; imposing biologically realistic constraints on interactions makes stable, diverse food webs substantially more likely [55]. Physical constraints further restrict the architectures available to ecological systems, with energetic requirements delimiting feasible trophic configurations [56]. These processes act at different levels and need not optimize the same quantity. Their combined effect can nevertheless favor architectures that gain the benefits of ecological integration without becoming so strongly coupled that responses lose their differentiation. High thermodynamic efficiency may therefore be an emergent signature of ecological assembly rather than its explicit objective.

The null-model results provide further insight into how this balance is encoded in ecological network architecture. Accounting for species-level degree heterogeneity brings the expected efficiency closer to empirical observations, suggesting that the coexistence of specialist and generalist species contributes to the balance between propagation and response diversity. Nevertheless, substantial positive deviations from configuration-model expectations remain, particularly in mutualistic networks. These deviations become considerably smaller when the empirical degree sequence is preserved exactly. However, this result should be interpreted cautiously, as stronger constraints on network randomization can restrict the space of possible configurations, potentially limiting the structural differences between empirical and randomized networks [48–50]. Thus, the contribution of higher-order organization cannot be established solely from the magnitude of these deviations. Nevertheless, the arrangement of interactions is known to have important ecological consequences. In mutualistic communities, nested and related structures can alter competition and enlarge the conditions compatible with coexistence [48, 57, 58], while modular and compartmentalized structures influence persistence and constrain the propagation of disturbances in trophic networks [59, 60]. Moreover, the relatively weak association between thermodynamic and global efficiency suggests that the propagation–response-diversity trade-off captures aspects of network organization beyond shortest-path accessibility. A shared link–species scaling regime therefore need not imply a shared detailed architecture: different ecological communities can realize the same broad connectivity constraint through different arrangements of interactions.

This perspective also places the classical complexity–stability problem within the broader question of how interactions accumulate with diversity. Random-matrix theory established relative sparsity as one way of preventing increasing diversity and interaction density from destabilizing large communities [26], and subsequent theory has connected link–species scaling more directly to robustness and local stability [28]. Stability provides one constraint on ecological connectivity, alongside network structure and environmental variation. Biologically constrained networks can sustain diverse stable communities well beyond the conditions predicted by unconstrained random matrices [55], while response diversity strongly influences temporal stability [38]. Recent work has further shown that structural heterogeneity can itself promote dynamical stability, with effects depending on the nature of ecological interactions [61]. The intermediate scaling found here is therefore better viewed as a broad architectural regime within which these dynamical mechanisms operate. The thermodynamic framework adds a complementary reason for relative sparsity: connectivity itself changes two collective properties at once, increasing integration across the network while reducing the independence of its responses.

Our interpretation has clear limitations. The networks analyzed here are binary, so interaction strengths, signs and functional forms—all central to ecological dynamics—are absent. Directed networks were symmetrized, meaning that diffusion describes pathways through the interaction backbone rather than the biological direction in which effects travel. Diffusion is itself a minimal representation of perturbation dynamics and cannot reproduce the nonlinear responses of ecological populations. Likewise, spectral response diversity should not be equated with response diversity measured from species-specific reactions to environmental change. Our perturbation experiment establishes local efficiency against random additions and removals of interactions, rather than optimality across the full space of possible network architectures, and the analytical stationary condition remains a finite-size mean-field consistency result. Variation in sampling effort, the temporal window in which the networks are observed, taxonomic resolution and network completeness may further contribute to the dispersion around the global scaling relationship. Although the consistency of the pattern across contrasting network types supports its generality, whether it extends to underrepresented interaction types and to networks integrating multiple interaction types remains to be established. In addition, these limitations prevent a more mechanistic understanding of how the observed scaling and efficiency emerge during ecological network assembly, pointing directly to the need for models that link interaction-level processes to the resulting system-level architecture.

Community-assembly and eco-evolutionary models in which interactions can be gained, lost and rewired could test whether intermediate link scaling and high thermodynamic efficiency emerge from species-level ecological and evolutionary processes, without explicit selection for network efficiency. Extending the framework to weighted, directed and signed interactions would clarify how interaction strength, asymmetry and sign reshape the propagation–response-diversity trade-off, while temporal networks could reveal how this balance changes during community assembly and following disturbance. Further progress will also require more comprehensive empirical data capturing interactions across entire ecological communities, their strengths, and the distinction between absent and undetected interactions. Together, these approaches could establish whether the trade-off identified here actively shapes ecological network assembly or emerges as a consequence of other constraints. Our findings suggest that the growth of ecological network complexity may reflect a general balance between increasing connectivity and preserving the diversity of possible responses to perturbations.

## Methods

### Data

The primary dataset used in this study consists of ecological interaction networks obtained from the Web of Life database and the Mangal.jl interface to the *Mangal* ecological network database. These repositories provide curated collections of ecological networks across multiple ecosystem types and interaction categories, including food webs, host–parasite systems, and mutualistic networks such as plant–pollinator and seed-dispersal communities. The geographic distribution of the networks included in the dataset is shown in Fig. 1a.

### Network preprocessing

In total, 1,801 interaction networks were initially compiled across different ecological interaction types. We verified that all networks were comprised of at least 3 species and contained at least one interaction for every species, as otherwise such systems do not represent meaningful ecological communities. We also excluded networks with zero variance in species degree. Inspection revealed that these were exclusively composed of disconnected pairs of interacting species. Such configurations lack pathways connecting different interacting pairs and fail to characterize the collective organization of ecological communities. We also removed duplicates networks contained in both datasets.

After applying these criteria, the dataset used for the structural scaling analysis comprised 1,533 ecological networks. For analyses involving thermodynamic efficiency and diffusion dynamics, we further restricted the dataset to communities with species richness *S >* 5, as the spectral properties of very small networks can produce degenerate diffusion modes. This resulted in a subset of 1,268 networks used in the efficiency analyses.

Because interaction data are reported using different conventions across datasets, all networks were converted to unweighted adjacency matrices where edges represent the presence or absence of interactions. For networks originally reported as directed (e.g., trophic interactions), adjacency matrices were symmetrized so that diffusion dynamics capture the existence of interaction pathways irrespective of direction. This representation is consistent with the diffusion framework adopted here, which focuses on the propagation of perturbations through the structural interaction network rather than on specific dynamical mechanisms associated with interaction directionality. Accordingly, the diffusion and efficiency analyses reported here should be interpreted as properties of the undirected structural interaction backbone rather than as direct representations of biologically directed interaction flows.

### Connectance

Connectance (*C*) quantifies the density of realized interactions relative to the total number of possible interactions in a network. Because the set of possible interactions depends on whether the community is represented as a unipartite or bipartite network, connectance was computed differently for these two classes.

### Unipartite networks

Food webs were treated as unipartite networks in which all species belong to a single set and interactions may occur between any pair of species. For an undirected network with *S* species and *L* realized interactions the total number of possible interactions is *L_max_* = *S*(*S* − 1)*/*2 (excluding self-interactions). Connectance was therefore computed as

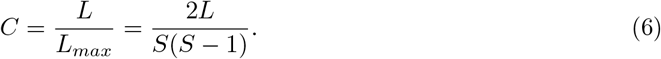

### Bipartite networks

Mutualistic systems (e.g., plant–pollinator and seed-dispersal networks) and other bipartite communities (e.g., host-parasite) were represented as networks composed of two distinct sets of species, such as plants and animals. Interactions are only allowed between species belonging to different sets. If the two sets contain *S*_1_ and *S*_2_ species, respectively, the total number of possible interactions is *L_max_* = *S*_1_*S*_2_. Connectance was therefore computed as

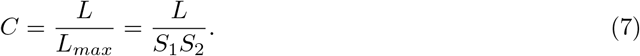

### Diffusion in ecological networks

We represent each ecological community as an undirected network of *S* species, with adjacency matrix **A**, where *A_ij_* = 1 if species *i* and *j* interact and *A_ij_* = 0 otherwise. The degree matrix **D** is diagonal, with *D_ii_* = *k_i_* = *_j_ A_ij_*, where *k_i_* is the number of interaction partners of species *i*. The graph Laplacian is then defined as

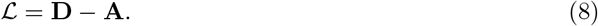

We use diffusion on this network as a minimal and general representation of how signals propagate through ecological interactions. Here, a signal represents a generic quantity or perturbation that can spread between species, without specifying its ecological nature or whether its effects are beneficial or detrimental. Diffusion does not assume a particular biological mechanism (e.g., predator–prey regulation or mutualistic feedback), but instead captures how the underlying interaction structure mediates propagation. This approach is motivated by the fact that the linearized responses of several nonlinear dynamical processes can be mapped, exactly or approximately under suitable conditions, into diffusion dynamics [42].

Let ***ψ****_τ_* describe the distribution of a signal across species at propagation scale *τ*. Assuming a uniform diffusion rate along each interaction, absorbed into *τ*, the dynamics are governed by [62]

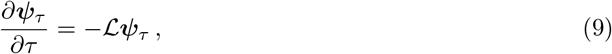

whose solution can be written as

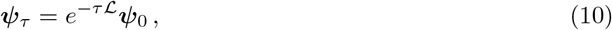

where ***ψ***_0_ is the initial signal distribution.

The diffusion operator *e*^−^*^τ^*^L^ determines how a signal initially localized at one species spreads throughout the network. In particular, its element (*e*^−^*^τ^*^L^)*_ji_* gives the fraction of a unit signal initially located at species *i* that reaches species *j* after propagation scale *τ*. The parameter *τ* therefore controls the extent of propagation through the interaction network: at small *τ*, signals remain largely localized around directly interacting species, whereas increasing *τ* allows them to reach more distant species through indirect pathways.

An intuitive understanding of how signals propagate through the network comes from the adjacency matrix **A**. A useful property of this matrix is that the element (**A***^n^*)*_ji_* counts the number of walks of length *n* connecting species *i* and *j*. For instance, **A**^2^ counts connections through an intermediate species, while higher powers account for increasingly long interaction pathways. Related matrix-power expansions have been used to quantify indirect evolutionary effects transmitted through successive interaction pathways in mutualistic networks [63, 64]. This property helps interpret the Taylor expansion of the diffusion operator,

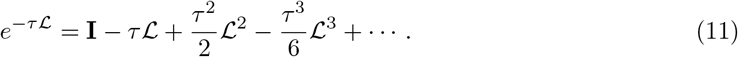

The zeroth-order term represents the initial state, before any propagation has occurred. Since −L = **A** − **D**, the first-order term allows signals to reach directly connected species. Higherorder terms progressively incorporate second, third, and more distant neighbors through indirect interaction pathways. Thus, the propagation scale *τ* controls the contribution of interactions at different distances, with larger values allowing signals to spread through increasingly distant parts of the network.

### Network density matrix

While the diffusion operator describes how a signal spreads from a given initial condition, we are interested in the collective response of the network to signals originating at different species. Indeed, the consequences of a local perturbation depend not only on how far it can propagate, but also on which species are affected and how their responses are related. By considering different possible origins, we can therefore characterize both the capacity of the network to distribute signals and the diversity of the resulting response patterns.

To do so, we represent each species *i* by a canonical row vector **v***_i_*, whose only non-zero element is at position *i*. A unit signal initially localized at that species is thus represented by **v***^t^*, and its subsequent distribution is given by *e*^−^*^τ^*^L^**v***^t^*. Following [42], we define the corresponding local propagator as

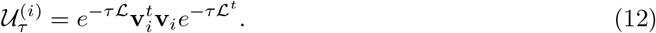

This matrix provides a description of how the network responds to a signal originating at species *i*. Its diagonal elements describe the distribution of signal energy (i.e., squared signal amplitudes) among species, whereas its off-diagonal elements encode correlations between pairs of species receiving the same signal. The local propagator therefore captures not only where a signal spreads, but also how the responses of different species are related.

Since signals may originate at any species, we next combine these local responses to obtain a statistical description of the entire network. In the absence of information about the relative likelihood of different origins, we assume that each species can be initially perturbed with equal probability, 1*/S*. Averaging the local propagators gives the statistical propagator,

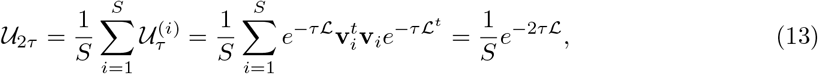

where we have used the symmetry of the Laplacian (L = L*^t^*), since we consider undirected networks, and Σ_*i*_ **v**^*t*^_*i*_ **v**_*i*_ = **I**. After rescaling the propagation parameter (2*τ* → *τ*), the statistical propagator takes the same form as the original diffusion operator, up to a factor 1*/S*. It thus summarizes the network’s possible responses to signals originating throughout the community, rather than its response to a particular perturbation.

The statistical propagator thus contains information about both the distribution of signal energy across the network and the correlations between species’ responses. Its trace, obtained by summing its diagonal elements, represents the total signal energy in the system. This provides a natural way to normalize the statistical propagator, allowing us to describe the relative distribution of signal energy independently of its total magnitude. Following [40–42], we define the network density matrix as

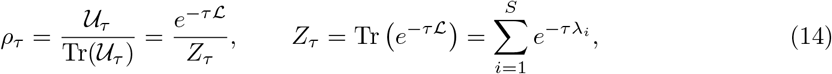

where {*λ_i_*}^*S*^_i=1_ are the eigenvalues of the graph Laplacian. The quantity *Z_τ_*, proportional to the trace of the statistical propagator, can be interpreted as a partition function within a statistical-mechanical framework, providing the normalization required for the density matrix to have unit trace.

The density matrix therefore retains the information contained in the statistical propagator, but expresses it in a probabilistic form. In particular, its diagonal elements represent the probability that a unit of signal energy is located at a given species, while its off-diagonal elements retain information about correlations between species’ responses. This normalized representation provides the basis for applying statistical-mechanical tools to characterize the network’s capacity to transport signals and the diversity of its possible responses, as described in the following subsection.

### Thermodynamic efficiency of signal propagation

The network density matrix allows us to characterize signal propagation and response diversity using concepts from statistical mechanics. In statistical mechanics, the properties of a system can be described through the relative contributions of its possible states, from which quantities such as free energy and entropy can be derived. Here, an analogous description is obtained from the diffusion modes of the network, which represent different collective patterns through which signals can spread and relax across species. These modes correspond to the eigenvectors of the graph Laplacian, while their eigenvalues determine how rapidly they decay. Modes with larger eigenvalues decay faster, whereas those with smaller eigenvalues remain relevant over longer propagation scales. The eigenvalues of the density matrix,

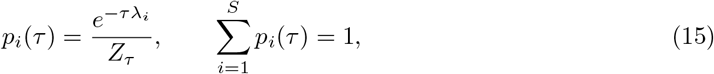

therefore quantify the relative contribution of each diffusion mode at propagation scale *τ*. This statistical representation provides the basis for defining two complementary quantities that characterize the network’s capacity to transport signals and the diversity of its possible responses [42].

The first is the network free energy,

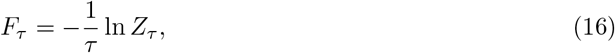

which quantifies the capacity of the network to transport signals among its components. To understand this interpretation, recall that the partition function *Z_τ_* is the trace of the diffusion operator, and therefore reflects the tendency of signals to be found at their original locations after propagation. Consequently, lower values of *Z_τ_* correspond to higher free energy and greater transport capacity.

The second quantity is the von Neumann entropy,

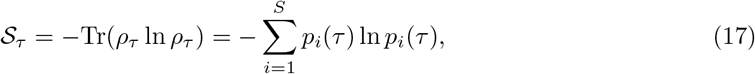

which quantifies the diversity of diffusion modes contributing to the network’s response. When many modes contribute with comparable weights, the network retains a greater diversity of possible response patterns and entropy is high. Conversely, when the response is dominated by a small number of collective modes, entropy is low. This quantity thus captures spectral response diversity, rather than variation in species abundances or species-specific responses to environmental change.

Ghavasieh and De Domenico [42] showed that, relative to an initial reference state *G*_0_ consisting of disconnected nodes, the introduction of interactions enhances signal propagation (increasing free energy) but reduces response diversity (decreasing entropy). These opposing effects can be quantified as a gain in free energy, *W_τ_* = *F_τ_* (*G*) − *F_τ_* (*G*_0_) ≥ 0, and an entropy-associated cost, *Q_τ_* = [S*_τ_* (*G*) − S*_τ_* (*G*_0_)]*/τ* ≤ 0, which mathematically resemble work and heat in thermodynamics, respectively. Their relative balance defines the thermodynamic efficiency,

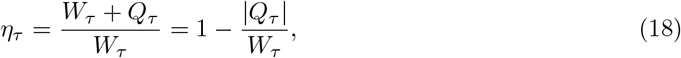

which ranges between zero and one for networks with *W_τ_ >* 0. High efficiency therefore characterizes networks that achieve substantial gains in signal propagation while minimizing the relative loss of response diversity.

### Optimal scaling from thermodynamic efficiency

The thermodynamic efficiency *η* cannot be obtained in closed form for arbitrary network topologies. Nevertheless, an approximate expression can be derived by combining a second-order expansion of the entropy and free energy with a mean-field treatment of the Laplacian spectrum [42]. Within this approximation, one assumes that the number of links scales with the number of nodes as *L* ∼ *cS^δ^*^+1^, where *c* and *δ* are treated as effective parameters characterizing network density. Under these assumptions, the efficiency can be approximated as

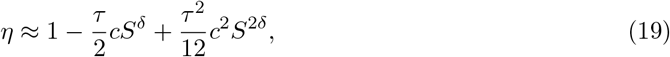

where *τ* denotes the diffusion time-scale.

Formally differentiating Eq. (19) with respect to *δ* yields

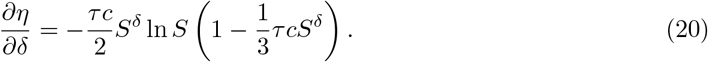

In the asymptotic limit *S* → ∞, this expression admits solutions corresponding either to decreasing connectivity with system size (*δ <* 0) or to sparse scaling (*δ* = 0) [42]. However, empirical ecological networks are finite, and the applicability of asymptotic optimality arguments is therefore limited. For finite *N*, Eq. (20) can be solved formally for the stationary point of the approximation, yielding

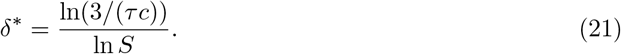

This expression should not be interpreted as defining a universal or self-consistent scaling exponent. Rather, it provides a heuristic indication of how the efficiency-maximizing balance between signal propagation and response diversity depends on system size, coupling strength, and dynamical time-scale within the adopted approximation. In particular, the dependence of *δ*^∗^ on *S* reflects the fact that the original scaling ansatz assumes a constant exponent, whereas the stationary condition is evaluated for finite systems. We therefore treat Eq. (21) as a consistency estimate rather than as a predictive scaling law. Any agreement between *γ_opt_* and the empirical exponent should therefore be interpreted as support for compatibility with the efficiency approximation, not as evidence of a uniquely identified generative mechanism.

In practice, we estimate the effective scaling of connectivity by fitting *L* = *cS^γ^*, with *γ* = *δ* + 1 to the whole dataset. Then, for each empirical network, the diffusion time *τ* is taken as the characteristic diffusion time-scale *τ*_diff_ = 1*/λ*_2_, where *λ*_2_ is the smallest non-zero eigenvalue of the Laplacian. These quantities are then used to evaluate Eq. (21) at the observed system size, allowing comparison between the empirically inferred scaling exponent and the value suggested by the efficiency approximation.

### Null network models

To determine whether the observed diffusion properties arise from generic structural features or from more specific ecological organization, we compared empirical networks with a hierarchy of null ensembles imposing progressively stronger structural constraints.

First, we generated Erdös–Rényi (ER) random networks preserving the number of nodes and the expected connectance of each empirical network. In these networks, interactions are placed randomly between pairs of species with equal probability. For bipartite networks, randomization was restricted to interactions between the two node sets, thereby preserving the bipartite structure. Second, we generated configuration-model (CM) networks accounting for the empirical degree heterogeneity. For unipartite networks, we generated configuration-model networks from the empirical degree sequence, subsequently removing self-loops and multiple edges to obtain simple graphs. For bipartite networks, we adopted the canonical bipartite configuration model (BiCM), motivated by probabilistic null models previously used in ecological networks [48] and formulated within a maximum-entropy framework [65, 66]. Thus, the model preserves species-level degree heterogeneity in expectation, while allowing individual degrees and the total number of interactions to fluctuate among randomized networks.

Finally, we used the Curveball (CB) algorithm as a more restrictive null model for bipartite networks [47]. This algorithm randomizes the incidence matrix through repeated pairwise trades while preserving its row and column sums exactly. Consequently, every species retains its empirical degree, and the total number of interactions remains unchanged. In contrast to the probabilistic BiCM, Curveball therefore explores networks with the observed degree sequence fixed in every realization. It is important to note that preserving the degree sequence exactly can severely restrict the space of admissible network configurations [48–50], so negligible deviations under Curveball do not necessarily imply that the degree sequence alone determines thermodynamic efficiency. Conversely, non-zero deviations indicate that rearranging interactions while preserving every species’ degree can change thermodynamic efficiency, revealing contributions from higher-order network organization.

For each empirical network and null ensemble, we generated 100 randomized networks and computed their thermodynamic efficiency. We quantified deviations from each null model as

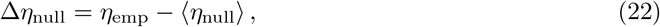

where the brackets denote the mean across randomized networks. Positive values indicate higher empirical efficiency than the corresponding null expectation. To quantify the prevalence of substantial efficiency gains across interaction types, we additionally calculated the fraction of networks satisfying Δ*η*_CM_ *>* 0.1.

### Interaction perturbation experiments

To assess the sensitivity of network efficiency to small structural perturbations, we constructed modified versions of each empirical network in which the total number of interactions was either increased or decreased by a given percentage *p*%. For interaction addition experiments, pairs of previously unconnected species were randomly selected and assigned new interactions until the desired increase in interaction number was reached. For interaction removal experiments, existing interactions were randomly selected and removed. For bipartite networks, interaction additions were restricted to pairs of nodes belonging to different partitions. Self-loops and duplicate edges were not allowed. For each network and perturbation type, an ensemble of 100 perturbed networks was produced for the subsequent analysis.

We quantified the effect of interaction perturbations by computing the difference in network efficiency between empirical and perturbed networks across a logarithmically spaced range of propagation scales, *τ* ∈ [10^−2^, 10^2^], and then averaging Δ*η* over *τ*. This provides a scale-averaged measure of whether empirical architectures are locally more efficient than nearby networks with slightly more or fewer interactions.

### Global efficiency

To quantify the topological efficiency of ecological interaction networks, we computed the standard global efficiency metric introduced in network science [67]. Global efficiency measures how efficiently information can be exchanged across a network through shortest paths.

For a network with *S* species, global efficiency is defined as

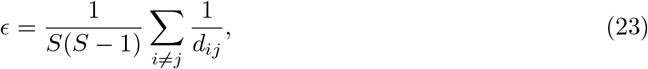

where *d_ij_* denotes the length of the shortest path between species *i* and *j*. If two species are not connected by any path, *d_ij_* = ∞ =⇒ *d*^−1^ = 0, so that efficiency is defined as zero. This metric captures the average inverse distance between all pairs of nodes and therefore increases when networks contain shorter paths or a higher degree of connectivity.

Global efficiency was computed for each empirical network using the implementation provided in the NetworkX Python library. We then compared global efficiency directly with thermodynamic efficiency, evaluated at each network’s characteristic diffusion scale *τ*_diff_ = 1*/λ*_2_, allowing us to assess whether networks exhibiting higher thermodynamic efficiency also tend to facilitate communication through shorter interaction pathways.

## Data availability

The original ecological interaction networks used in this study are available from the Web of Life database and the Mangal ecological network database. The processed dataset generated for this study, including the networks analysed in GraphML format and the derived structural and thermodynamic-efficiency metrics, is publicly available on Zenodo [68].

## Code availability

All code used to process the networks, compute thermodynamic efficiency, generate null models, perform perturbation analyses and reproduce figures is available in a public GitHub repository [69] and archived on Zenodo [68].

## Author contributions

A.G.R., D.O., J.B. and M.G. conceived and developed the study. A.G.R. compiled the datasets, developed the computational methods, performed the analyses, prepared the figures and wrote the original manuscript. All authors contributed to interpreting the results, revising the manuscript and approving its final version.

## Competing interests

The authors declare no competing interests.

## Acknowledgments

We thank Rob Salguero-Gómez, Pablo Almaraz and György Barabás for feedback in previous versions of this work. AGR acknowledges financial support from grant JDC2024-053275-I, funded by MICIU/AEI/10.13039/501100011033 and FSE+. AGR and MG acknowledge financial support from the Spanish Ministerio de Ciencia e Innovación/AEI and EU-FEDER (PID2021-124731NB-I00 and PIE 202430E234). AGR, DO and MG acknowledge financial support from the Spanish Ministerio de Ciencia e Innovacíon/AEI (PID2025-174810NB-I00).

## Notes

https://zenodo.org/records/23800150

https://github.com/agimenezromero/Ecological-networks-balance-connectivity-with-flexibility-to-perturbations

### Competing Interest Statement

The authors have declared no competing interest.

### Summary of Updates

This version has been substantially revised to focus on the intermediate link species scaling observed across ecological interaction networks and its possible explanation through a thermodynamic trade off between signal propagation and response diversity. We expanded the empirical analyses to examine scaling across individual interaction types and separately for bipartite and unipartite networks. We introduced a new conceptual figure, revised the null model comparisons, and extended the network perturbation analyses. We also clarified the ecological interpretation of thermodynamic efficiency and its possible emergence through community assembly, while explicitly acknowledging alternative explanations and limitations. The complexity stability analysis has been moved to the Supplementary Information and reframed as a theoretical benchmark. Finally, we updated the dataset to account for cross database duplicates, resulting in 1533 ecological networks, and revised the figures, methods, supplementary analyses, and supporting code accordingly.

