## Supplementary Information for "Ecological networks balance connectivity with flexibility to perturbations"

October 30, 2026

### 1 Scaling of connectance and species richness from stability constraints

#### 1.1 The Generalized Lotka-Volterra framework

We model the dynamics of an ecological community of $S$ interacting species using the Generalized Lotka-Volterra (GLV),

$$\frac{dn_i}{dt} = \dot{n}_i = n_i \left( r_i + \sum_{j=1}^{S} A_{ij} n_j \right) , \tag{1}$$

where $n_i$ is the abundance of species $i$, $r_i$ is its intrinsic growth rate, and $A_{ij}$ are the elements of the interaction matrix $\mathbf{A}$, representing the effect of species $j$ on the per-capita growth rate of species $i$.

A fixed point (equilibrium) $\mathbf{n}^*$ occurs when the rate of change for all species is zero, $\frac{dn_i}{dt} = 0$. For any species with non-zero abundance ($n_i^* \neq 0$), this implies,

$$r_i + \sum_{j=1}^{S} A_{ij} n_j^* = 0 \,, \tag{2}$$

which solution can be found as,

$$\mathbf{r} + \mathbf{A}\mathbf{n}^* = 0 \implies \mathbf{n}^* = -\mathbf{A}^{-1}\mathbf{r} \,. \tag{3}$$

A fixed point is defined as feasible if and only if all equilibrium abundances are strictly positive:

$$\mathbf{n}^* > \mathbf{0} \,. \tag{4}$$

This condition links the network structure ($\mathbf{A}$) and the external environment ($\mathbf{r}$) to the possibility of coexistence.

#### 1.2 Local asymptotic stability

To evaluate the stability of a feasible equilibrium $\mathbf{n}^*$ against infinitesimal perturbations, we perform a linear stability analysis. The dynamics of such perturbations are given by

$$\boldsymbol{n}(t) = \boldsymbol{n}^* + \boldsymbol{\xi}(t) \;, \tag{5}$$

where $\xi$ is the magnitude of the perturbation to the species' densities, considered small.

We obtain a linearised approximation of the system by taking a Taylor expansion of the system of ODEs around the equilibrium and discarding higher order terms:

$$\frac{\mathrm{d}\boldsymbol{\xi}(t)}{\mathrm{d}t} = \boldsymbol{M}\boldsymbol{\xi} \,, \tag{6}$$

where $\boldsymbol{M}$ is the Jacobian matrix of the system evaluated at the equilibrium, also known as the community matrix, given by

$$m_{ij} = \left.\frac{\partial \dot{n}_i}{\partial n_j}\right|_{\boldsymbol{n}^*} \,, \tag{7}$$

where $m_{ij}$ are the elements of the matrix $M$, that is, the linearisation of the ODEs at equilibrium densities, defined by the partial derivatives of the species growth rate with respect to the species density evaluated at the equilibrium.

In the case of Eq. (1), this specifically reads:

$$m_{ij} = n_i^* A_{ij} = -\left(\sum_{k=1}^{S} \left(A^{-1}\right)_{ik} r_k\right) A_{ij} \,. \tag{8}$$

The solution of the linearised system (6) is given by

$$\xi_i(t) = \sum_{j=1}^{S} \mathfrak{C}_{ij} e^{\lambda_j t} \,, \tag{9}$$

where $\mathfrak{C}_{ij}$ are constants that depend on the initial conditions and $\lambda_j$ are the eigenvalues of the community matrix $M$.

Thus, the stability of the system is determined by the sign of the real parts of the eigenvalues of the community matrix. If all eigenvalues have negative real parts, all terms in Eq. (9) decay to zero and the system is locally asymptotically stable. However, if there exists even one eigenvalue with a positive real part, the corresponding term in Eq. (9) grows exponentially, driving the population away from the equilibrium and rendering the system locally unstable.

##### 1.3 May's complexity stability relationship

Following the framework established by May [1], we assume $\mathbf{A}$ is a random matrix with certain constraints. The intra-specific interactions (i.e., the diagonal terms $A_{ii}$) are fixed to a given negative value, resembling self-regulation arising due to negative density-dependence. The inter-specific interactions (i.e., the off-diagonal terms $A_{ij}$) are non-zero with probability $C$ (the connectance). Non-zero entries are drawn from a distribution with mean zero and variance $\sigma^2$.

According to the circular law for random matrices, the eigenvalues of $\mathbf{A}$ are uniformly distributed in a disk in the complex plane with radius $R = \sigma\sqrt{SC}$ [2, 3]. Then, for the fixed point to be locally asymptotically stable, the real part of all eigenvalues of $\mathbf{M}$ must be negative (Eq. (9)). This requires the radius of the eigenvalue distribution of $\mathbf{A}$ to be smaller than the magnitude of the diagonal coefficients, $A_{ii} = d$,

$$\sigma\sqrt{SC} < d \,. \tag{10}$$

This is the so called complexity-stability relationship. May assumed that all diagonal coefficients were fixed to the same number, $A_{ii} = d = 1$. However, later extensions have shown that $d$ can be understood as the mean of the diagonal coefficients, $d = \mathbb{E}(A_{ii})$ [3].

Isolating the connectance $C$, we define the theoretical upper boundary for stability, $C_{max}$,

$$C_{max} = \frac{1}{S}\left(\frac{d}{\sigma}\right)^2 \implies C \propto S^{-1} \,. \tag{11}$$

This derivation establishes that for a system to remain stable as species richness $S$ grows, the connectance must decrease as the inverse of $S$, provided that the ratio of the variance of the inter-specific interactions, $\sigma$, and the mean of the intra-specific interactions, $d$, do not scale with species richness, $S$.

#### 1.4 Accounting for feasibility

While local stability addresses small perturbations, a biologically meaningful community must also be feasible—meaning all species maintain positive equilibrium abundances ($\mathbf{n}^* > 0$). The conditions under which a community is both feasible and globally stable have been recently studied [4, 5].

A sufficient condition ensuring that any feasible interior equilibrium is globally asymptotically stable is that the interaction matrix $\mathbf{A}$ be negative definite [5–8]. A condition for a matrix $\mathbf{M}$ to be negative definite is determined by the largest eigenvalue of the symmetric part of the matrix, $(\mathbf{M} + \mathbf{M}^T)/2$ [6]. For the case of random matrices, this can be expressed in terms of the moments of the elements [9]

$$-d + \max\left\{SE_1, -E_1 + \sqrt{2S\,(1+E_c)E_2}\right\} < 0\,, \tag{12}$$

where $E_1$, $E_2$ and $E_c$ are given by [5],

$$E_1 = \frac{1}{S(S-1)} \sum_{i=1}^{S} \sum_{j \neq i} A_{ij} \tag{13}$$

$$E_2 = \sqrt{\frac{1}{S(S-1)\sum_{j \neq i}(A_{ij})^2 - E_1^2}} \tag{14}$$

$$E_c = \frac{1}{E_2^2}\left(\frac{1}{S(S-1)} \sum_{i=1}^{S} \sum_{j \neq i} A_{ij}A_{ji} - E_1^2\right) \tag{15}$$

For the random matrix ensemble described above $E_1 = E_c = 0$ and $E_2 = \sqrt{\sigma^2 C}$, so that the expression simplifies to

$$-d + \max\left\{0, \sqrt{2S}\sqrt{\sigma^2 C}\right\} < 0 \tag{16}$$

This leads to the criterion,

$$-d + \sigma\sqrt{2SC} < 0 \tag{17}$$

Rearranging for $C$ yields the maximum connectance compatible with this sufficient condition for global stability of a feasible equilibrium,

$$C_{max} = \left(\frac{d}{\sigma}\right)^2 \frac{1}{2S} \implies C \propto S^{-1} \tag{18}$$

Remarkably, despite the different mathematical requirements of global stability and feasibility compared to local asymptotic stability, the functional dependence of $C$ on $S$ remains identical. The difference lies in the constant coefficient, with the sufficient condition for global stability of a feasible equilibrium imposing a more restrictive bound (by a factor of $1/2$) on the same inverse scaling law.

#### 1.5 Specific interaction types

In real ecological communities, the effect of species $j$ on $i$ is often related to the effect of $i$ on $j$. We define the correlation $\rho$ between pairs of off-diagonal elements as

$$\rho = \frac{\mathrm{cov}(M_{ij}, M_{ji})}{\sigma^2} \tag{19}$$

where $\sigma^2$ is the variance of the non-zero off-diagonal elements. According to the Elliptical Law [2, 10], the eigenvalues of such a matrix (with connectance $C$ and diagonal $-d$) are distributed within an ellipse centered at $-d$. The horizontal axis (along the real line) has a length of $R = \sigma\sqrt{SC}(1+\rho)$.

For the system to be locally asymptotically stable, the rightmost edge of this ellipse must remain to the left of the imaginary axis ($R < d$). This leads to a generalized stability criterion,

$$\sigma\sqrt{SC}(1+\rho) < d \tag{20}$$

Using this criterion, the stability-complexity limit can be obtained by communities rendering specific interaction types [11],

$$\begin{aligned}
&\textbf{Mutualistic:} && \gamma_m = \sigma C(S-1)\sqrt{\tfrac{2}{\pi}} < d \\
&\textbf{Competitive:} && \gamma_c = \sigma\left\{\sqrt{SC}\left(1+\tfrac{2-2C}{\pi-2C}\right)\sqrt{\tfrac{\pi-2C}{\pi}} + C\sqrt{\tfrac{2}{\pi}}\right\} < d \\
&\textbf{Predator-prey:} && \gamma_p = \frac{\pi-2}{\pi}\sigma\sqrt{SC} < d \\
&\textbf{Mixture:} && \gamma_x = \frac{\pi+2}{\pi}\sigma\sqrt{SC} < d
\end{aligned} \tag{21}$$

By isolating the maximum connectance $C_{max}$ in each case, and considering $S \gg 1$, we can observe how the structural constraints of the community are shaped by the nature of the links,

- **Mutualistic communities:**

$$C_{max} = \frac{d}{\sigma S}\sqrt{\frac{\pi}{2}} \implies C \propto S^{-1} \tag{22}$$

- **Predator-prey communities:**

$$C_{max} = \frac{1}{S}\left(\frac{d\pi}{\sigma(\pi-2)}\right)^2 \implies C \propto S^{-1} \tag{23}$$

- **Competitive communities:** Although the expression for $\gamma_c$ is algebraically transcendental, we consider the limit for large $S$. In this regime, the term proportional to $\sqrt{SC}$ dominates, as $C$ is indeed bounded between 0 an 1. This leads to the approximate limit:

$$C_{max} \approx \frac{1}{S}\left(\frac{d}{\sigma}\right)^2 \implies C \propto S^{-1} \tag{24}$$

- **Mixture of competition and mutualism:**

$$C_{max} = \frac{1}{S}\left(\frac{d\pi}{\sigma(\pi+2)}\right)^2 \implies C \propto S^{-1} \tag{25}$$

These results show that the $C \sim S^{-1}$ scaling appears across all fundamental ecological interaction types. While the specific interaction type dictates the "efficiency" of the architecture—shifting the coefficient and allowing for varying levels of absolute density—the functional dependence on species richness remains invariant. This convergence reinforces the view that the complexity-stability paradox is not a contradiction, but a reflection of general dynamical constrains on network connectivity.

#### 1.6 Stability in meta-communities

The impact of spatial structure and dispersal on community stability has been investigated through the lens of meta-community theory, which accounts for the arrangement of populations across multiple patches. By considering a meta-community of $n$ patches, Gravel et al. [12] derived a modified complexity-stability relationship that incorporates the effects of spatial scale and synchrony:

$$\sigma\sqrt{\frac{C(S-1)\left[1+(n-1)\rho\right]}{n}} < d\,, \tag{26}$$

where $n$ represents the number of patches and $\rho$ characterizes the spatial correlation of species' interactions or environmental responses across the network.

To determine the architectural limit imposed by spatial structure, we isolate the maximum connectance $C_{max}$ that satisfies the stability criterion:

$$C_{max} = \left(\frac{d}{\sigma}\right)^2 \frac{n}{[1 + (n-1)\rho]\,(S-1)}\,. \tag{27}$$

For large communities ($S \gg 1$), this expression converges to the same fundamental scaling law:

$$C_{max} \approx \frac{1}{S}\left(\frac{d}{\sigma}\right)^2 \frac{n}{1 + (n-1)\rho} \implies C \propto S^{-1}\,. \tag{28}$$

This result confirms that the inclusion of spatial dimensions—while potentially shifting the absolute stability threshold depending on the number of patches ($n$) and their degree of correlation ($\rho$)—does not alter the underlying scaling. Whether an ecological system is modeled as a single local community or a spatially distributed meta-community, the inverse relationship between richness and connectance remains an essential constraint for maintaining stability.

#### 1.7 Synthesis of scaling laws

The theoretical derivations presented above demonstrate a remarkable convergence across different ecological frameworks. Whether considering local asymptotic stability, global stability and feasibility, specific interaction types, or meta-community dynamics, the maximum allowable connectance $C_{max}$ for a persistent community consistently follows an inverse scaling with species richness $S$.

We can summarize these findings into a common theoretical benchmark:

$$C_{max} = \kappa S^{-1}\,, \tag{29}$$

where $\kappa$ is a constant determined by the specific biological and structural assumptions of the model. The table below summarizes the theoretical values of $\kappa$ derived from the various frameworks:

**Table 1: Summary of complexity-stability scaling constants across different theoretical models.**

| Framework | Constraint | Effective Limit ($\kappa$) |
|---|---|---|
| May (1972) [1] | Local Stability | $(d/\sigma)^2$ |
| Grilli et al. (2017) [5] | Global stability conditional on feasibility | $\frac{1}{2}(d/\sigma)^2$ |
| Allesina & Tang (2012) [11] | Predator-Prey | $\left(\frac{d\pi}{\sigma(\pi-2)}\right)^2$ |
| Allesina & Tang (2012) [11] | Mutualism | $\frac{d}{\sigma}\sqrt{\frac{\pi}{2}}$ |
| Allesina & Tang (2012) [11] | Mixture | $\left(\frac{d\pi}{\sigma(\pi+2)}\right)^2$ |
| Gravel et al. (2016) [12] | Meta-communities | $(d/\sigma)^2\frac{n}{1+(n-1)\rho}$ |

While $\kappa$ varies—reflecting how different interaction types (e.g., predator–prey) or spatial structures (e.g., metacommunities) modify the conditions for stability—the inverse dependence $C_{\max} \propto S^{-1}$ recurs across these theoretical frameworks. This convergence suggests a potentially general architectural constraint: as communities become more diverse, the maximum interaction density compatible with the stability criteria considered decreases inversely with species richness. However, the resulting boundary depends on the assumptions of each model and should therefore be regarded as a theoretical benchmark rather than a universal limit for empirical ecological communities.

#### 2 Empirical scaling laws

The theoretical derivations in Section 1 provide a classical reference for how connectance can scale with species richness under a range of random-matrix stability assumptions. Despite differences among the particular formulations, they frequently yield the asymptotic benchmark $C_{\max} \propto S^{-1}$. As emphasized above, this scaling depends on restrictive assumptions regarding interaction strengths, self-regulation, and network structure, and we therefore do not interpret it as a direct prediction of the stability of the empirical communities analysed here. Instead, we use it as a simple theoretical reference against which to characterize the observed diversity–connectance relationship.

Across the empirical dataset, connectance declines systematically with species richness according to a robust power law, $C \sim S^{-0.65}$ (95% CI $[-0.67, -0.64]$, $R^2 = 0.78$, $p < 0.001$; Fig. 1b). Interaction-type-specific fits yielded the same qualitative decline across food webs, host–parasite networks, pollination networks, seed-dispersal systems, plant–ant networks, and anemone–fish associations (Fig. 4). Although the magnitude of the exponent varied among interaction types, the estimates were broadly consistent within their 95% confidence intervals. The same qualitative relationship was recovered when bipartite and unipartite networks were analysed separately (Fig. 5). The interaction classes nevertheless differed substantially in their typical size and structural organization, including degree heterogeneity, modularity, and core structure (Table 3).

The empirical exponent is substantially shallower than the $-1$ scaling obtained from the classical theoretical benchmarks (Fig. 1a,b). This discrepancy indicates that some of the quantities treated as richness-independent in these simplified derivations must themselves vary systematically with community size, or that additional structural effects become increasingly important as communities grow. In the generic expression

$$C_{\max} = \kappa S^{-1}, \tag{1}$$

the coefficient $\kappa$ collects the dependence on interaction-strength statistics, self-regulation, and the particular structural assumptions of the model (Table 1). If, as a simple illustration, this coefficient varied with richness as $\kappa(S) \propto S^x$, the resulting scaling would become $C_{\max} \propto S^{-(1-x)}$; the empirical exponent $-0.65$ would correspond formally to $x \simeq 0.35$. Within May's simplest formulation, where $\kappa = (d/\sigma)^2$, such a dependence could arise from systematic changes in self-regulation $d$, in the variance of inter-specific interaction strengths $\sigma^2$, or in both. More generally, the departure need not be reducible to a richness-dependent prefactor: non-random network organization can alter the community-matrix spectrum through degree heterogeneity and isolated structural eigenvalues, potentially changing which spectral component constrains stability [13, 14]. The observed scaling therefore suggests that ecological communities depart systematically from the size-independent assumptions of the classical random-matrix benchmark, although the binary network data analyzed here do not allow us to distinguish among these possible mechanisms.

A long-standing alternative explanation for declining connectance is the constant-degree hypothesis [15, 16]. Under this hypothesis, each species maintains an approximately fixed number of interaction partners as community size increases. The total number of interactions should therefore increase linearly with richness, $L \propto S$. For unipartite networks, $C = 2L/[S(S-1)] \propto L/S^2$ for large $S$, so that $L \propto S$ implies $C \propto S^{-1}$. For bipartite networks, $C = L/(S_1 S_2)$, leading to the same asymptotic dependence when the relative sizes of the two guilds do not change systematically with total richness.

Our results reject a strict constant-degree explanation. If species maintained a fixed number of interaction partners, average node degree would remain independent of richness. Instead, average degree increases systematically with system size, following $\langle k \rangle \sim S^{0.43}$ ($R^2 = 0.54$, $p < 0.001$; Fig. 1c), and the same qualitative result holds within interaction types (Fig. 8). Thus, species in larger communities interact, on average, with progressively more partners, although this increase is not sufficient to maintain constant connectance. The degree distributions are also broad and heterogeneous, with highly connected species occurring alongside many species with fewer interactions (Fig. 7), further excluding a strict constant-degree architecture.

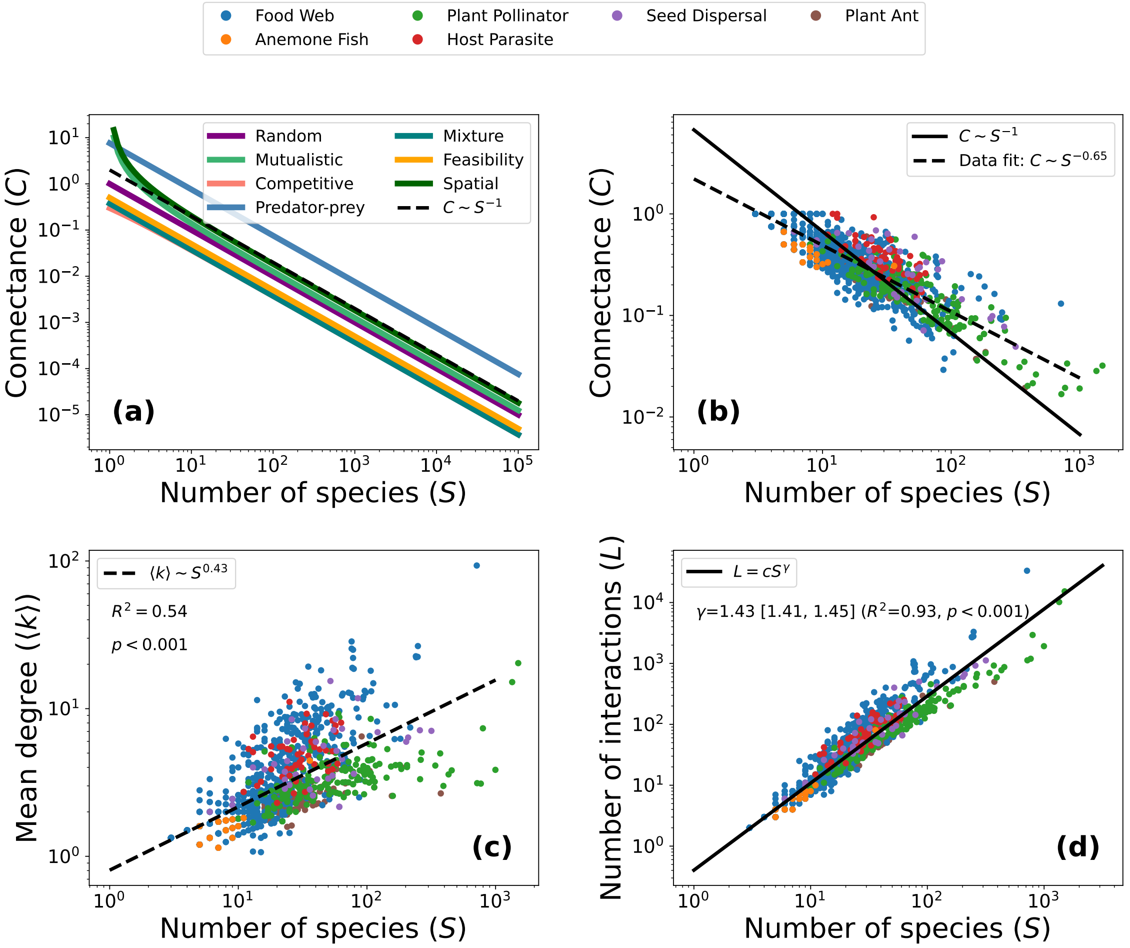


**Supplementary Figure 1: Diversity–connectance and interaction scaling in ecological communities.** (**a**) Classical theoretical benchmarks in the connectance–diversity ($C$–$S$) parameter space. Lines show $C_{\max} \propto S^{-1}$ relationships obtained under different random-matrix stability formulations, including local stability [1, 11], a sufficient condition for global stability conditional on feasibility [5], and metacommunity dynamics [12]. These relationships are shown as theoretical references rather than universal stability boundaries for empirical communities (derivations and assumptions in Section 1). Values of $C > 1$ are formal extrapolations with no ecological meaning. (**b**) Empirical scaling of connectance with species richness for $\sim$ 1500 ecological networks. Data points are color-coded by interaction type (see legend). The dashed line indicates the best fit for the global dataset, $C \sim S^{-0.65}$ with $\alpha = -0.65$ (95% CI: [-0.67, -0.64]), $R^2 = 0.78$, $p < 0.001$), which follows a smaller slope than the random-system theoretical baseline of $S^{-1}$ (solid black line). (**c**) Scaling of the average node degree ($\langle k \rangle$) with species richness. Contrary to the constant-degree hypothesis ($L \sim S$), the average number of interactions per species increases significantly with community size ($\langle k \rangle \sim S^{0.43}$, $R^2 = 0.54$, $p < 0.001$), indicating that the observed sparsity is unlikely to arise from a strict individual-level constant-degree constraint alone. (d) Scaling of the total number of interactions with species richness, $L \sim S^{\gamma}$ ($\gamma = 1.43$ [1.41, 1.45], $R^2 = 0.93$, $p < 0.001$).

These relationships define an intermediate scaling regime. Connectance decreases with richness ($C \sim S^{-0.65}$), while average degree increases ($\langle k \rangle \sim S^{0.43}$), implying that the total number of interactions grows superlinearly with community size, $L \sim S^{1.43}$ (Fig. 1d). Ecological networks therefore occupy a regime between the linear scaling associated with constant degree ($L \sim S$) and the quadratic scaling of a fully connected network ($L \sim S^2$). Classical complexity–stability arguments provide one reason why unrestricted growth of connectance may be dynamically consequential, but they do not explain why empirical networks consistently occupy this particular intermediate scaling regime. Explaining the observed link-scaling relationship therefore requires considering constraints on network organization beyond the classical random-matrix benchmark.

#### Link–species scaling

To test whether the global link–species relationship arises from pooling structurally distinct ecological networks, we fitted

$$L \sim S^{\gamma}$$

separately within each interaction class. Interaction richness increased superlinearly with species richness in every case (Fig. 3). The estimated exponents ranged from $\gamma = 1.22$ for plant–pollinator networks to $\gamma = 1.63$ for food webs, with plant–ant ($\gamma = 1.27$), host–parasite ($\gamma = 1.28$), seed-dispersal ($\gamma = 1.28$), and anemone–fish ($\gamma = 1.60$) networks falling between these values. In all interaction classes, the 95% confidence intervals excluded both $\gamma = 1$ and $\gamma = 2$. Thus, the intermediate link–species regime is recovered independently within networks spanning different interaction types, although the rate at which interactions accumulate with diversity differs among them.

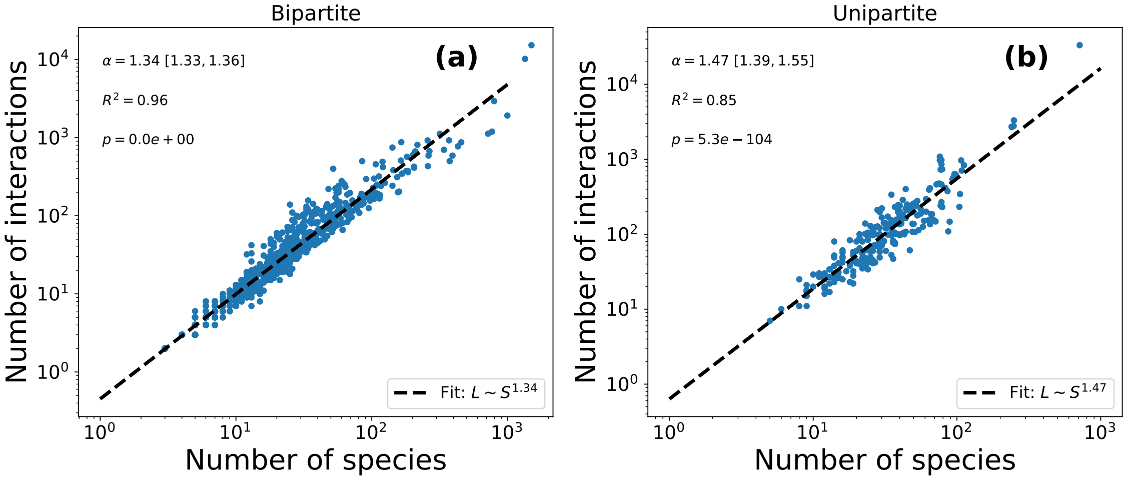


**Supplementary Figure 2: Link–species scaling in bipartite and unipartite ecological networks.** Relationship between species richness ($S$) and interaction richness ($L$) for **(a)** bipartite and **(b)** unipartite ecological networks. Dashed lines show the best-fitting power-law relationship, $L \sim S^{\gamma}$, fitted separately for each network class; estimated exponents, 95% confidence intervals, coefficients of determination, and statistical significance are reported in each panel. Interaction richness increased superlinearly with species richness in both network representations, with bipartite networks following $L \sim S^{1.34}$ and unipartite networks following $L \sim S^{1.47}$. In both cases, the estimated exponents lie between the limiting expectations of constant degree ($\gamma = 1$) and constant connectance ($\gamma = 2$), showing that the intermediate link–species scaling regime is robust to differences in network representation, although the rate of interaction accumulation differs between bipartite and unipartite systems.

We next asked whether this variation was associated with the broad structural representation of the networks. When analyzed separately, bipartite networks followed $L \sim S^{1.34}$ (95% CI: [1.33, 1.36]), whereas unipartite networks scaled more steeply, as $L \sim S^{1.47}$ (95% CI: [1.39, 1.55]; Fig. 2). Both classes therefore remain within the same superlinear but sub-quadratic regime, while exhibiting quantitatively different scaling exponents.

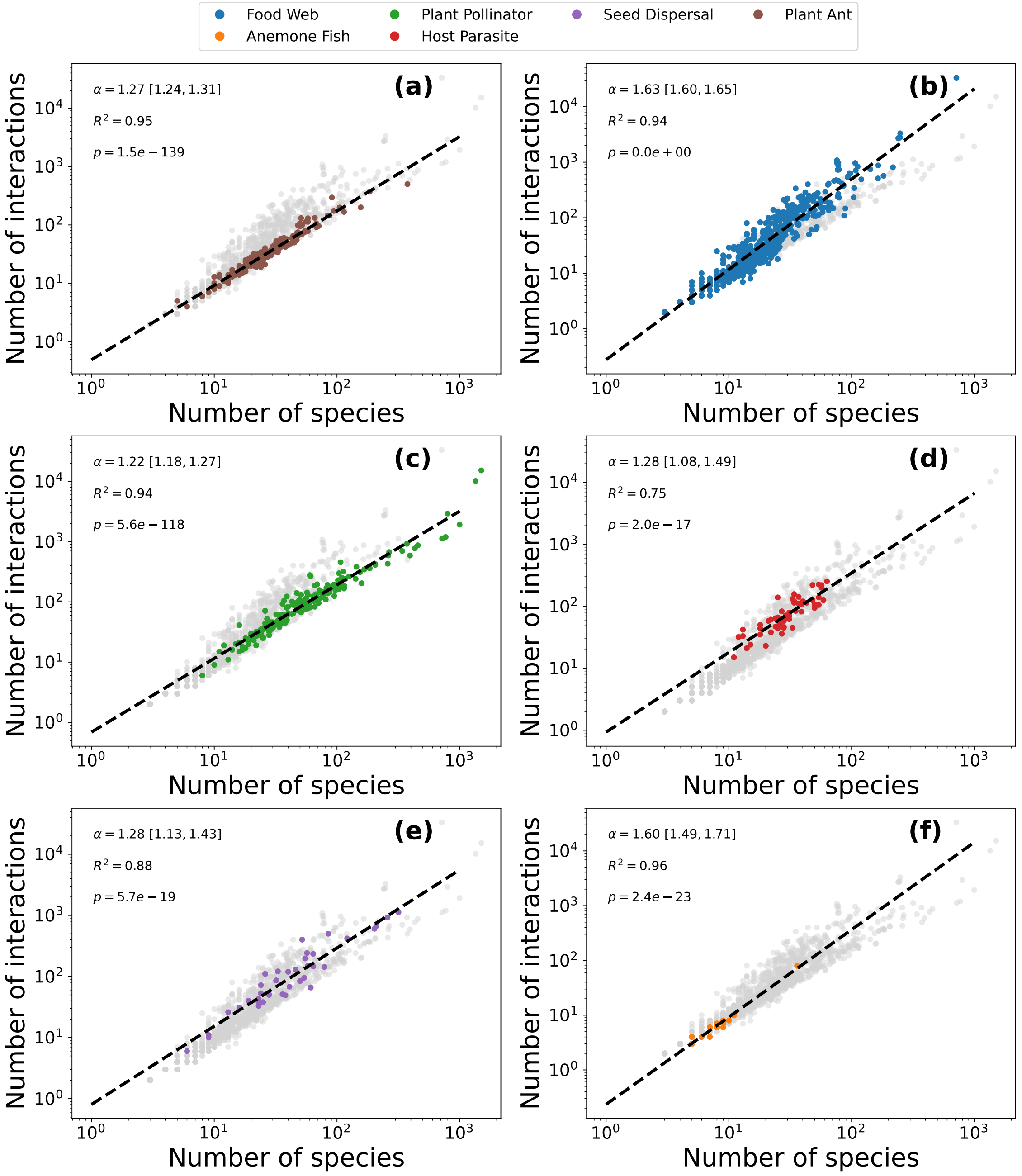


**Supplementary Figure 3: Link–species scaling across ecological interaction types.** Relationship between species richness ($S$) and interaction richness ($L$) for **(a)** plant–ant, **(b)** trophic, **(c)** plant–pollinator, **(d)** host–parasite, **(e)** seed-dispersal, and **(f)** anemone–fish networks. Colored points denote networks of the focal interaction type and grey points show the remaining networks for reference. Dashed lines indicate the best-fitting power-law relationship, $L \sim S^{\gamma}$, fitted separately within each interaction class; estimated exponents, 95% confidence intervals, coefficients of determination, and statistical significance are reported in each panel. Interaction richness increases superlinearly with species richness in every interaction class, with exponents ranging from $\gamma = 1.22$ for plant–pollinator networks to $\gamma = 1.63$ for food webs. All confidence intervals fall between the limiting expectations of constant degree ($\gamma = 1$) and constant connectance ($\gamma = 2$), showing that different interaction networks independently occupy an intermediate link–species scaling regime while differing in the rate at which interactions accumulate with diversity.

Together, these analyses show that the global $L \sim S^{1.43}$ relationship is not an artifact of combining networks with different characteristic sizes or architectures. The robust feature is not a universal value of $\gamma$, but an intermediate regime in which interaction richness grows faster than species richness while remaining far below quadratic growth. Interaction type and network representation determine where ecological networks fall within this regime.

##### Comparison to other studies

Previous studies have reported different relationships between interaction richness and species richness, particularly in food webs and plant–animal mutualistic networks (Table 2). Although several studies identified intermediate scaling exponents, their estimates vary with taxonomic resolution, network sampling, interaction definitions, and statistical methodology. Moreover, previous comparative analyses have focused predominantly on trophic networks, with fewer studies considering other interaction types. Our analysis extends these comparisons to a broader range of ecological interactions and shows that intermediate scaling consistently emerges across mutualistic and antagonistic networks of different types, although the corresponding exponents differ among interaction types.

**Table 2:** Previously reported link–species scaling relationships and comparison with the present study. Unless specified otherwise, $\gamma$ denotes the exponent in $L \propto S^{\gamma}$, with $L$ the number of observed interactions and $S$ the number of species. Estimates are just descriptive comparisons: sampling design, taxonomic resolution, estimation procedures, and the definition of an interaction differ among studies.

| Study | Interaction type | Networks | $\gamma$ | Interpretation and comparability |
|---|---|---|---|---|
| Cohen et al.[17,18] | Food webs | 113 | 1.356 | Reported for taxonomically aggregated community food webs; not considered significantly different from a linear fit. |
| Martinez[19] | Food webs | 175 | $\approx 2$ | Constant-connectance interpretation; estimated scaling depends strongly on web size and taxonomic resolution. |
| Schmid-Araya[18] | Stream food webs | 22 | $1.299 \pm 0.07$ | Higher-resolution stream food webs; slope differs significantly from both 1 and 2. |
| Bascompte[20] | Pollination and seed dispersal | 52 | 1.139 | Combined analysis of plant–animal mutualistic networks, rather than separate interaction-type estimates. |
| Banašek-Richter[21] | Food webs (topological) | — | 1.62 | Each realized trophic link counted once; directly analogous to binary link counts, subject to data and fitting differences. |
| Banašek-Richter[21] | Food webs (quantitative) | — | 1.47 | Quantitative interaction complexity $L_q$, not the same response variable as binary link richness. |
| MacDonald[22] | Food webs | 255 | $1.70 \pm 0.043$ | Power-law fitted using a negative-binomial likelihood; a different, constrained model was preferred. |
| This study | Plant–ant | 207 | 1.27 $[1.24, 1.31]$ | 95% CI; Fig. 3(a). |
| This study | Food webs | 1009 | 1.63 $[1.60, 1.65]$ | 95% CI; Fig. 3(b). |
| This study | Plant–pollinator | 190 | 1.22 $[1.18, 1.27]$ | 95% CI; Fig. 3(c). |
| This study | Host–parasite | 55 | 1.28 $[1.08, 1.49]$ | 95% CI; Fig. 3(d). |
| This study | Seed dispersal | 40 | 1.28 $[1.13, 1.43]$ | 95% CI; Fig. 3(e). |
| This study | Anemone–fish | 32 | 1.60 $[1.49, 1.71]$ | 95% CI; Fig. 3(f). |
| This study | All networks | 1,533 | 1.43 $[1.41, 1.45]$ | 95% CI; Fig. 1(d). |

#### Diversity-connectance scaling

Because the number of structurally possible interactions differs among network representations, we next examined how this variation appears after normalizing interaction richness by the corresponding interaction space, through connectance. We examined the scaling within each interaction type separately to verify that the global relationship previously reported does not arise from mixing structurally distinct systems.

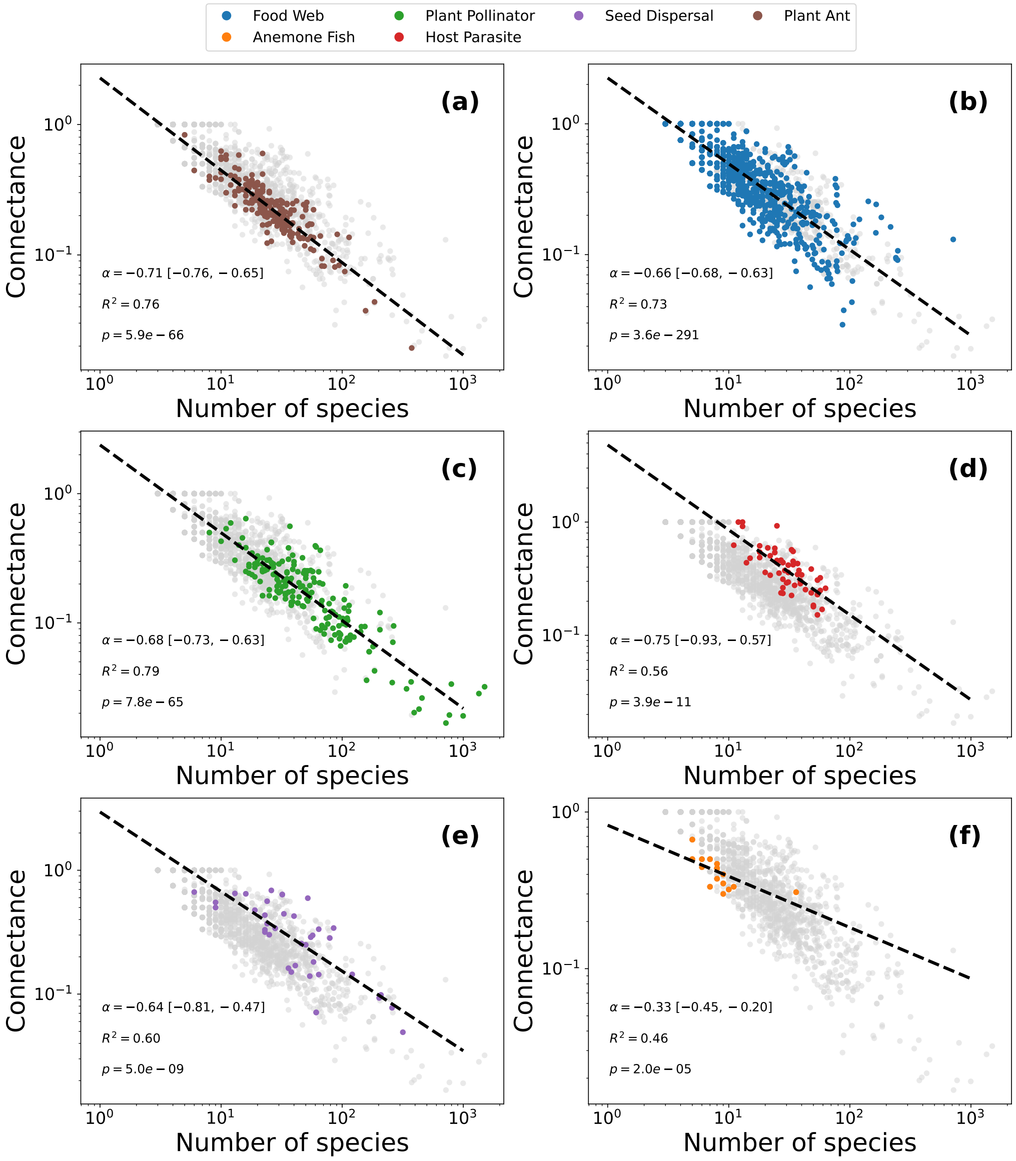


**Supplementary Figure 4: Scaling of connectance with species richness across interaction types.** Relationship between species richness ($S$) and connectance ($C$) for each interaction type analyzed in this study: food webs, plant–pollinator networks, seed-dispersal networks, plant–ant interactions, anemone–fish symbioses, and host–parasite systems. In all cases a significant negative power-law relationship is observed, indicating that interaction density declines systematically as communities become more diverse. Dashed lines show the best power-law fit $C \sim S^{-\alpha}$ for each interaction class, with reported coefficients and statistical significance. Despite differences in ecological mechanisms and network representation, the scaling exponents remain broadly consistent across interaction types.

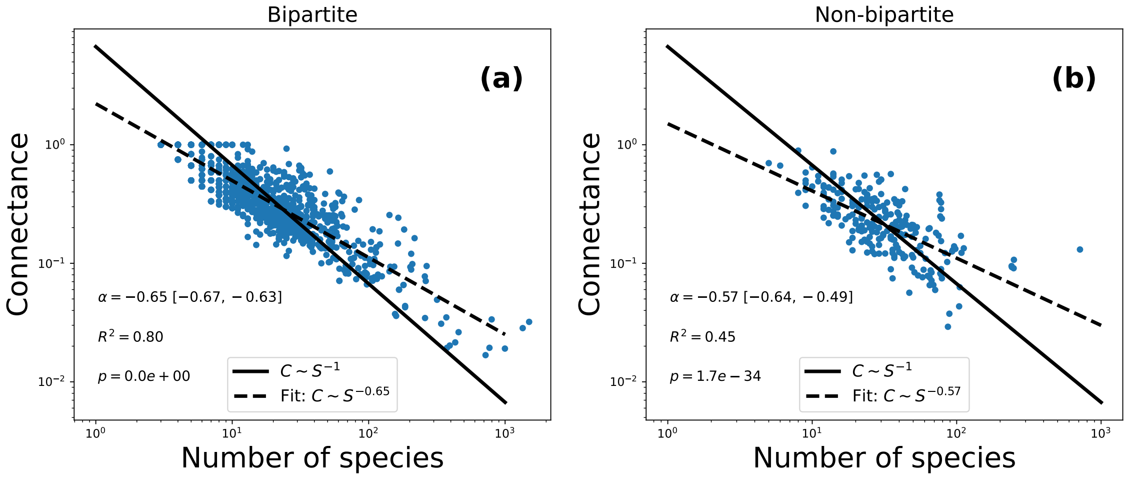


**Supplementary Figure 5: Scaling of connectance with species richness comparing bipartite with unipartite graphs.** Relationship between species richness ($S$) and connectance ($C$) for bipartite **(a)** and unipartite **(b)** networks. In both cases a significant negative power-law relationship is observed, indicating that interaction density declines systematically as communities become more diverse. Dashed lines show the best power-law fit $C \sim S^{-\alpha}$ for each class, with reported coefficients and statistical significance. Despite differences in ecological mechanisms and network representation, the scaling exponents remain broadly consistent across interaction types.

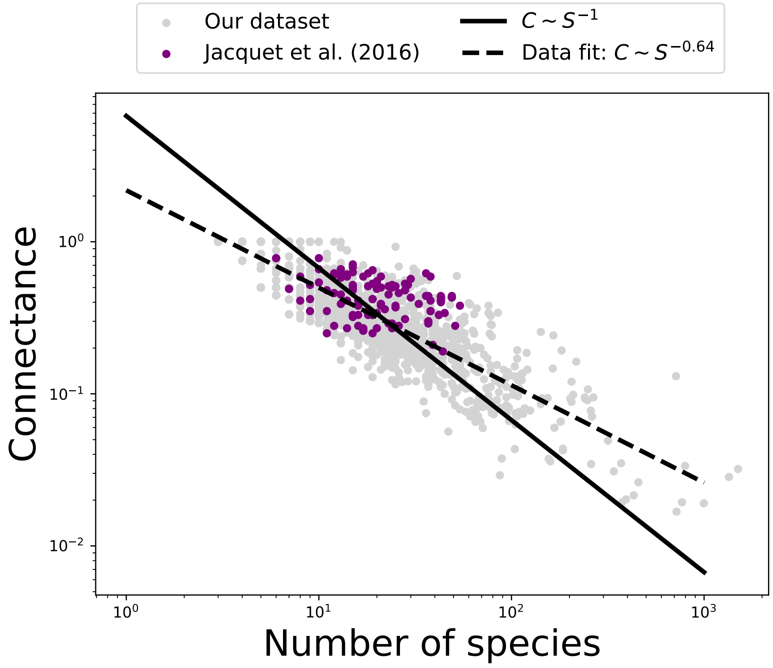


**Supplementary Figure 6: Comparison between the global dataset and the Ecopath food webs analyzed by Jacquet et al. (2016).** Connectance–richness ($C$–$S$) relationship for the ecological communities analyzed in this study (grey points) together with the 116 quantitative food webs examined by Jacquet et al. [23] (purple points). The solid black line represents the theoretical complexity-stability $C \propto S^{-1}$ derived from random matrix theory, while the dashed line indicates the best-fit scaling for our compiled dataset + Jaquet's data ($C \sim S^{-0.64}$). The Jacquet et al. networks fall within the variability of the broader dataset and follow the same structural trend, indicating that their reported absence of a complexity–stability relationship does not reflect a different structural regime in the $(S, C)$ parameter space.

Across all interaction types, connectance declines systematically with species richness following a power-law relation of the form $C \sim S^{-\alpha}$. Although the precise exponent varies slightly among interaction classes, the negative scaling is consistently observed across both antagonistic and mutualistic systems, as well as across unipartite and bipartite network representations (Figs. 4 and 5). This robustness indicates that the decrease of interaction density with diversity is a general architectural property of ecological communities rather than a feature specific to a particular interaction type.

To compare our structural scaling results with previous empirical work on the complexity–stability relationship, we projected the communities analysed by Jacquet et al. [23] onto the same species-richness–connectance ((S,C)) space used in the main analysis. The Jacquet et al. communities fall within the same broad ((S,C)) envelope as the interaction networks analysed here (Fig. 6). In particular, they are consistent with the general tendency for connectance to decline with species richness and occupy the region expected for the size of the ecological communities represented.

#### Average degree and degree distribution

The decline of connectance with species richness could arise trivially if species maintained an approximately constant number of interaction partners as communities become larger [15, 16]. To test this possibility, we examined the scaling of mean degree with species richness separately for each interaction type. Across interaction classes, mean degree generally increased with species richness (Fig. 8), indicating that larger communities are not simply formed by adding species with a fixed number of partners. Instead, interaction richness increases with community size, although not fast enough to maintain constant connectance.

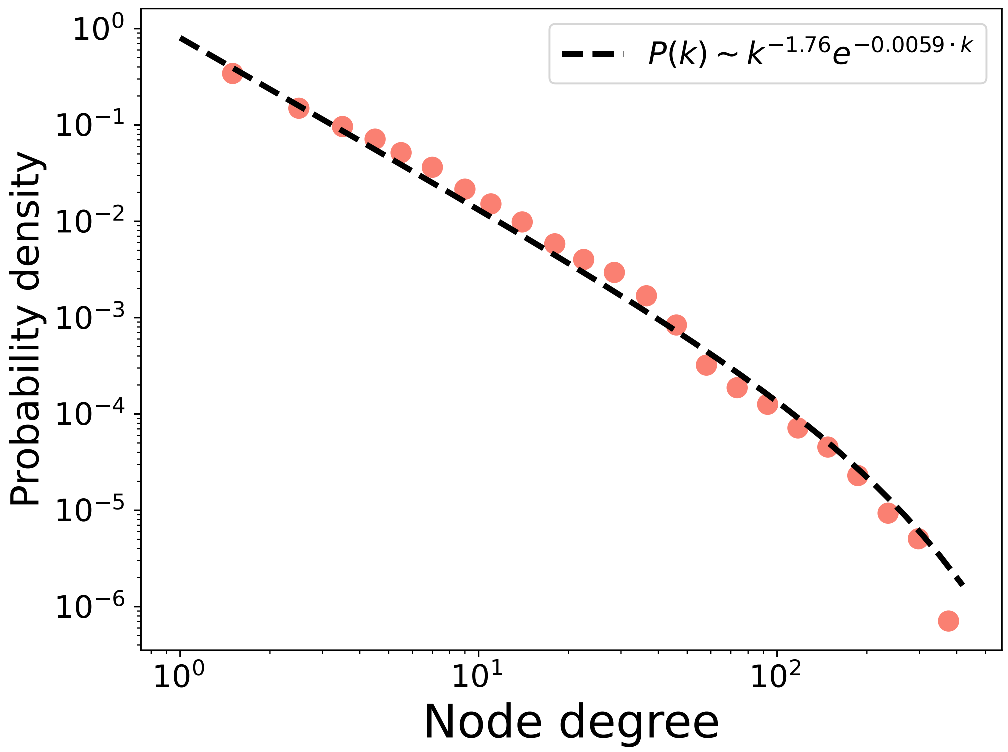


**Supplementary Figure 7: Degree distribution across the empirical ecological networks.** Probability distribution of node degree $P(k)$ aggregated across all networks in the dataset. The distribution exhibits a heavy-tailed form well described by a truncated power-law, $P(k) \sim k^{-1.76} e^{-0.0059 \cdot k}$, indicating the presence of highly connected hub species within larger communities. Such heterogeneity in interaction partners is incompatible with the constant-degree hypothesis and supports the conclusion that ecological networks exhibit strongly heterogeneous connectivity patterns.

We further examined the full degree distributions of the empirical networks. The degree distribution were broad and right-skewed, with many weakly connected species and a smaller number of highly connected species (Fig. 7). This heterogeneous organization is inconsistent with a strict constant-degree architecture and supports the interpretation that ecological networks occupy an intermediate structural regime: communities become progressively more interaction-rich as species richness increases, while remaining sparse relative to the number of possible interactions.

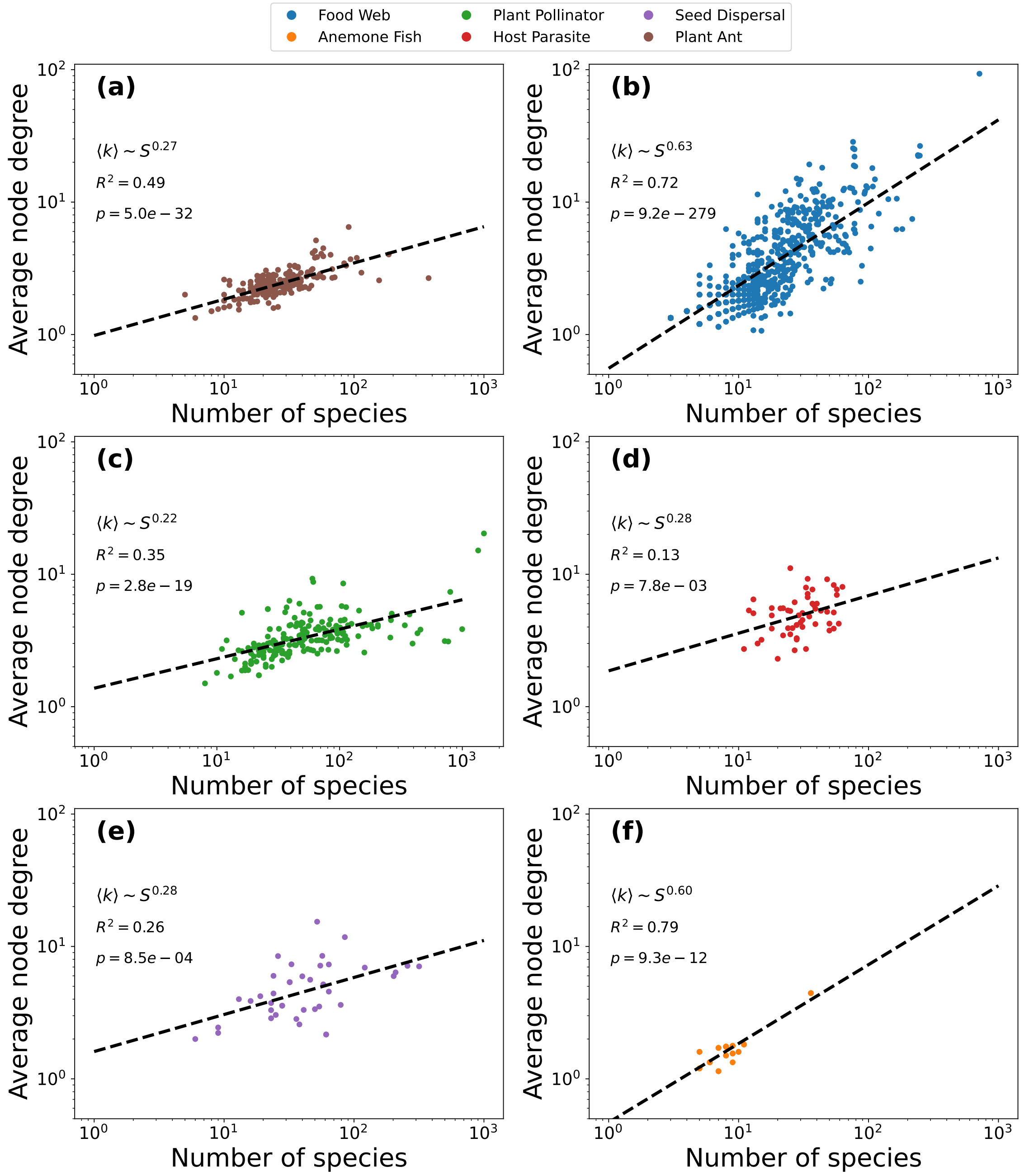


**Supplementary Figure 8: Scaling of average node degree with species richness across interaction types.** Relationship between species richness ($S$) and the average node degree ($\langle k \rangle$) computed for each ecological network, shown separately for **(a)** plant–ant, **(b)** food webs, **(c)** plant–pollinator, **(d)** host–parasite, **(e)** seed-dispersal, and **(f)** anemone–fish communities. Dashed lines indicate power-law fits of the form $\langle k \rangle \sim S^{\beta}$ (exponent, $R^2$, and $p$-value reported in each panel). Across all interaction types, $\langle k \rangle$ increases with richness, indicating that the observed decrease in connectance with $S$ is not explained by a constant-degree constraint.

#### 2.1 Robustness to the bounded nature of connectance

Because connectance is bounded to $0 < C \leq 1$, we additionally tested whether the diversity–connectance relationship was sensitive to the log–log representation used in the main analysis. We thus fitted a linear relationship between $\mathrm{logit}(C) = \log[C/(1-C)]$ and $\log S$.

The negative relationship remained strong under this alternative representation (Fig. 9), with $\mathrm{logit}(C) = -0.84 \log S + 1.76$ ($R^2 = 0.64$, $p < 0.001$). Thus, the decline of connectance with species richness is robust to explicitly accounting for the bounded nature of connectance.

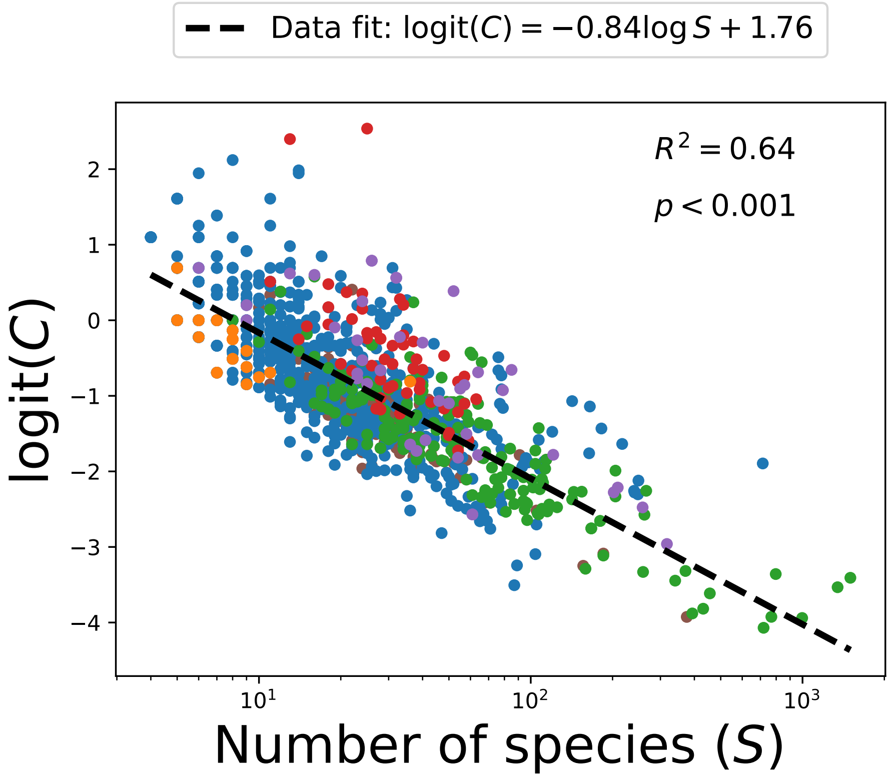


**Supplementary Figure 9: Diversity–connectance relationship under a logit transformation.** Relationship between species richness ($S$) and $\mathrm{logit}(C) = \log[C/(1-C)]$ across empirical ecological networks. The dashed line shows the linear fit $\mathrm{logit}(C) = -0.84 \log S + 1.76$ ($R^2 = 0.64$, $p < 0.001$). The persistence of the negative relationship under this alternative transformation indicates that the observed decline of connectance with species richness is not an artefact of representing the bounded variable $C$ on a logarithmic scale.

### 3 Structural characteristics across interaction types

The ecological network classes included in the dataset differed substantially in their typical size and structural organization (Table 3). To characterize this heterogeneity independently of the scaling analyses, we computed a set of basic structural descriptors for each network and report their median and interquartile range within each interaction class.

Plant–pollinator networks were typically larger and less connected than food webs and anemone–fish networks, consistent with the overall decline in connectance with species richness. Host–parasite networks showed relatively high mean degree and comparatively low modularity, suggesting a more connected and less compartmentalized architecture. Plant–ant and plant–pollinator networks exhibited higher modularity and greater relative degree heterogeneity. Thus, the common decline of connectance with species richness reported across interaction types occurs despite substantial differences in their underlying network architecture.

**Table 3:** Structural characteristics of the empirical ecological networks by interaction type. Values are reported as median [first quartile, third quartile] across networks within each interaction class. $n$ is the number of networks, $S$ species richness, $L$ the number of interactions, $C$ connectance, $\langle k \rangle$ the mean node degree, $\mathrm{CV}_k$ the coefficient of variation of node degree, and $Q$ modularity. Connectance was calculated relative to the set of possible interactions for each network representation, using the corresponding unipartite or bipartite definition.

| Interaction type | $n$ | $S$ | $L$ | $C$ | $\langle k \rangle$ | $\mathrm{CV}_k$ | $Q$ |
|---|---|---|---|---|---|---|---|
| Anemone–fish | 32 | 8 [7, 9] | 7 [4, 7.25] | 0.44 [0.33, 0.48] | 1.66 [1.46, 1.75] | 0.38 [0.33, 0.46] | 0.39 [0.35, 0.46] |
| Food web | 1009 | 10 [5, 20] | 9 [4, 34] | 0.50 [0.30, 1.00] | 1.83 [1.56, 3.47] | 0.58 [0.43, 0.77] | 0.23 [0.00, 0.36] |
| Host–parasite | 55 | 30 [24, 39] | 70 [45.5, 117] | 0.37 [0.28, 0.48] | 5.08 [3.90, 6.00] | 0.74 [0.64, 0.89] | 0.23 [0.16, 0.31] |
| Plant–ant | 207 | 25 [19, 34] | 30 [20, 45] | 0.24 [0.18, 0.30] | 2.38 [2.09, 2.75] | 0.89 [0.76, 1.05] | 0.45 [0.38, 0.50] |
| Plant–pollinator | 190 | 47 [27, 91] | 83 [39, 173] | 0.18 [0.11, 0.26] | 3.20 [2.68, 4.06] | 0.95 [0.78, 1.20] | 0.42 [0.36, 0.48] |
| Seed dispersal | 40 | 38 [24, 61] | 86 [49, 150] | 0.33 [0.16, 0.50] | 4.42 [3.32, 6.93] | 0.85 [0.63, 1.03] | 0.27 [0.19, 0.36] |

### 4 Network efficiency

#### 4.1 Efficiency curves for different interaction types

To characterize how perturbations propagate through ecological interaction networks, we compute the thermodynamic efficiency $\eta(\tau)$ introduced in the main text for each empirical network. This quantity captures the trade-off between the ability of the network to propagate information (e.g., energy) across species and the diversity of independent response modes available to the community.

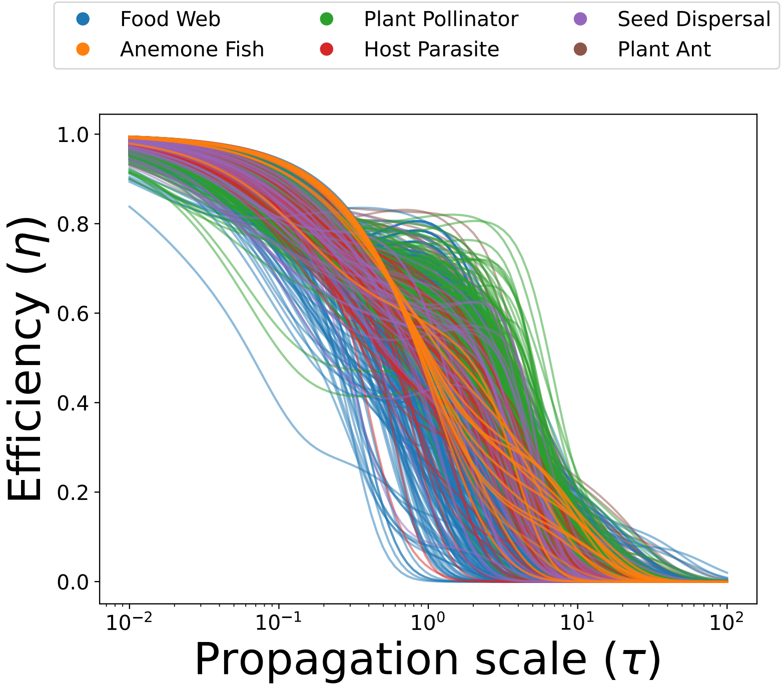


**Supplementary Figure 10: Efficiency profiles for individual ecological networks.** Thermodynamic efficiency $\eta(\tau)$ computed for individual networks across the dataset, grouped by interaction type. Each curve corresponds to a single empirical network. Efficiency is close to unity at short diffusion times, when perturbations remain localized, and decreases as $\tau$ increases and diffusion integrates larger portions of the network. The consistent qualitative behavior across interaction types indicates that ecological networks share a common dynamical signature in terms of information propagation.

Efficiency is evaluated as a function of the diffusion scale $\tau$ for all networks in the dataset. Fig. 10 shows the efficiency profiles for individual communities grouped by interaction type. These curves illustrate the characteristic behavior discussed in the main text: efficiency is close to unity at small diffusion times, when perturbations remain localized, and decreases progressively as diffusion integrates the network.

#### 4.2 Null models

To assess which structural properties contribute to thermodynamic efficiency, we compared empirical ecological networks with three null ensembles imposing different constraints on network architecture.Erdös–Rényi (ER) networks preserve network size and expected connectance but randomize interactions independently of species identity. Configuration-model (CM) networks additionally account for species-level degree heterogeneity. For bipartite networks, we used a probabilistic configuration model in which the expected degree of each species matches its empirical degree, while individual degrees can vary across realizations (see Methods). Finally, the Curveball (CB) algorithm provides a more restrictive comparison for bipartite networks by preserving the degree of every species exactly.

The comparison revealed substantial differences between empirical networks and both ER and CM ensembles (Figs. 11 and 12). ER networks generally exhibited lower thermodynamic efficiency, particularly at intermediate propagation scales, indicating that network size and expected connectance alone do not reproduce the empirical efficiency profiles. Accounting for species-level degree heterogeneity through the CM brought the null-model profiles closer to the empirical ones, but substantial deviations remained across interaction types. These differences are particularly evident in the distributions of efficiency deviations, which show predominantly positive values relative to both null ensembles, with especially pronounced deviations in plant–pollinator networks.

We further examined whether these deviations persisted when the empirical degree sequence was preserved exactly. Unlike the probabilistic CM, which allows species degrees and the total number of interactions to fluctuate among realizations, Curveball preserves both the degree of every species and the total number of interactions. Under these more restrictive constraints, efficiency deviations were substantially reduced and generally concentrated around zero (Fig. 13). Nevertheless, positive deviations remained in a considerable fraction of networks, indicating that some empirical architectures exhibit higher efficiency than the corresponding fixed-degree null expectation.

Together, these results highlight the importance of species-level connectivity in determining the thermodynamic efficiency of ecological networks, while also revealing contributions from the specific arrangement of interactions. Efficiency deviations relative to probabilistic CM networks become substantially smaller under Curveball, although this result should be interpreted cautiously: preserving the degree sequence exactly can severely restrict the space of possible configurations, so small or negligible deviations do not necessarily imply that degree heterogeneity alone determines efficiency. Conversely, the persistence of non-zero deviations under Curveball demonstrates that alternative arrangements of interactions with exactly the same species degrees can exhibit different thermodynamic efficiencies. Thus, even within the restricted space of degree-preserving configurations, higher-order network organization can influence the balance between propagation and response diversity.

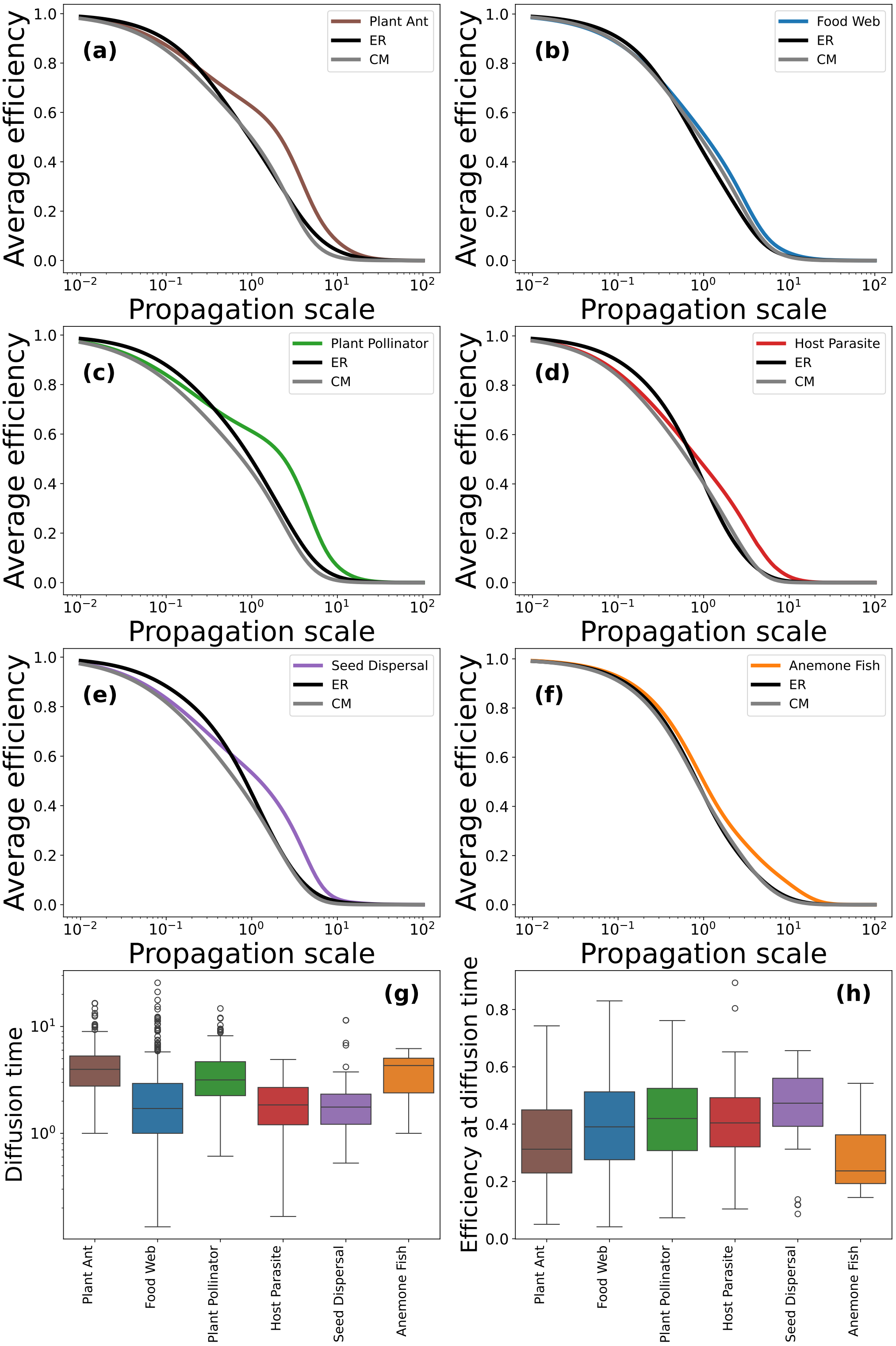


**Supplementary Figure 11: Comparison of empirical efficiency profiles with null network ensembles.** Thermodynamic efficiency $\eta(\tau)$ for empirical ecological networks (solid lines) compared with Erdös–Rényi (ER) and configuration-model (CM) null networks. ER networks preserve network size and expected connectance, whereas CM networks additionally account for empirical degree heterogeneity. Across interaction types, ER networks generally exhibit lower efficiency at intermediate propagation scales, while CM networks more closely reproduce the empirical profiles, although substantial differences remain. The bottom panels show the distributions of characteristic diffusion times, $\tau_{\text{diff}}$, and thermodynamic efficiency evaluated at these scales across interaction types.

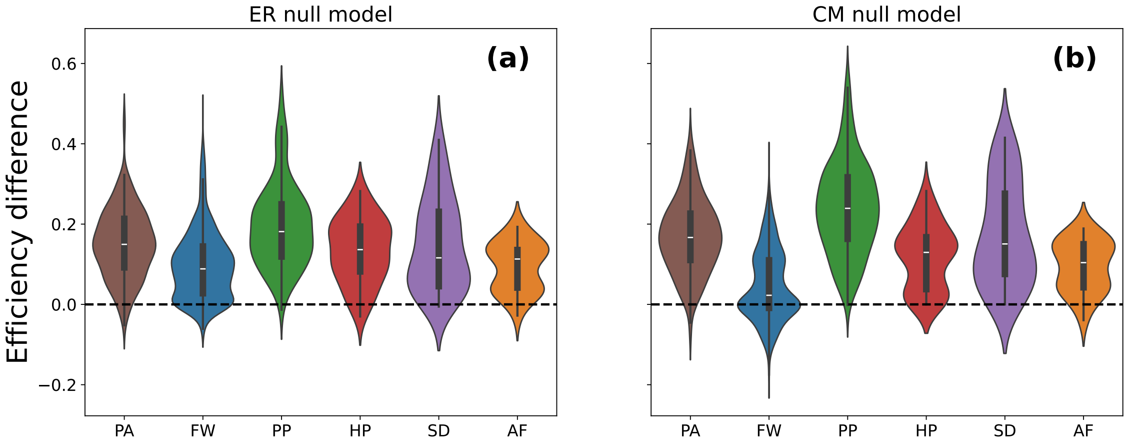


**Supplementary Figure 12: Thermodynamic efficiency deviations from ER and CM null models.** Distributions of differences in thermodynamic efficiency between empirical networks and **(a)**Erdös–Rényi (ER) and **(b)** configuration-model (CM) null ensembles, evaluated at the characteristic diffusion scale $\tau_{\text{diff}} = 1/\lambda_2$. Differences are defined as $\Delta\eta_{\text{null}} = \eta_{\text{emp}} - \langle\eta_{\text{null}}\rangle$, where brackets denote the mean across null-model realizations. Positive values indicate higher empirical efficiency than the corresponding null expectation. Substantial positive deviations are observed relative to both null models, particularly in plant–pollinator networks, indicating that expected connectance and degree heterogeneity alone do not fully account for the observed efficiency.

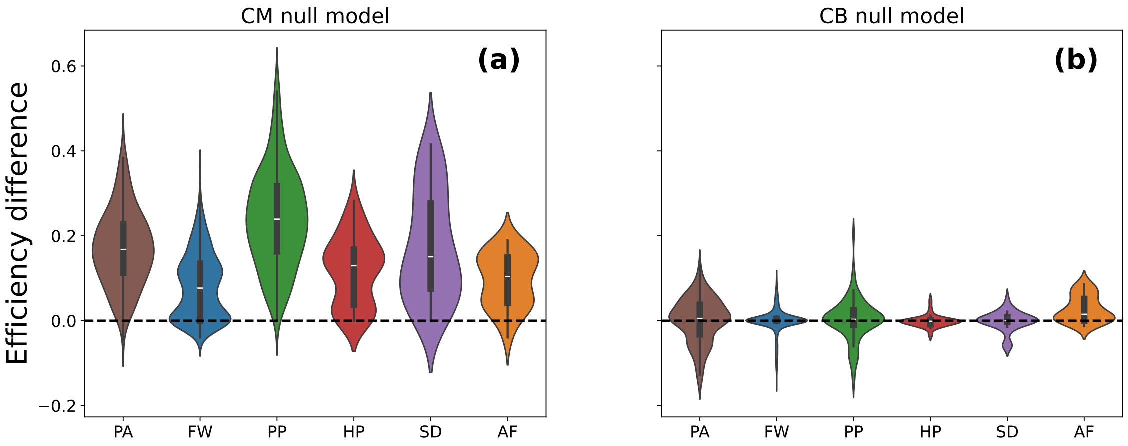


**Supplementary Figure 13: Comparison of efficiency deviations under probabilistic and fixed-degree null models.** Distributions of thermodynamic efficiency deviations relative to **(a)** configuration-model (CM) and **(b)** Curveball (CB) null ensembles. For bipartite networks, CM preserves the empirical degree of each species in expectation, while CB preserves the degree sequence and total number of interactions exactly. Positive deviations indicate higher empirical efficiency than the corresponding null expectation. Deviations are substantially reduced under CB and generally concentrated around zero, although positive deviations persist in several interaction classes. These results highlight the importance of the realized degree sequence while suggesting possible additional contributions from higher-order network organization.

#### 4.3 Perturbation experiments

To test whether the perturbation analysis depended on the arbitrary choice of a (1%) link modification, we repeated the analysis across a broader range of perturbation magnitudes. Specifically, for each empirical network we randomly added or removed (1%) to (10%) of its empirical number of links and quantified the resulting scale-averaged change in network efficiency, $\Delta\eta$, across

$\tau \in [10^{-2}, 10^2]$. Both link additions and removals produced consistently negative values of $\Delta\eta$ across the full perturbation range (Fig. 14), indicating that the empirical networks are locally more efficient than networks with either slightly higher or slightly lower link density. The effect was asymmetric, with link removals generally producing stronger reductions in $\eta$ than link additions, suggesting that empirical architectures may be particularly sensitive to link loss while still showing reduced efficiency under additional random coupling.

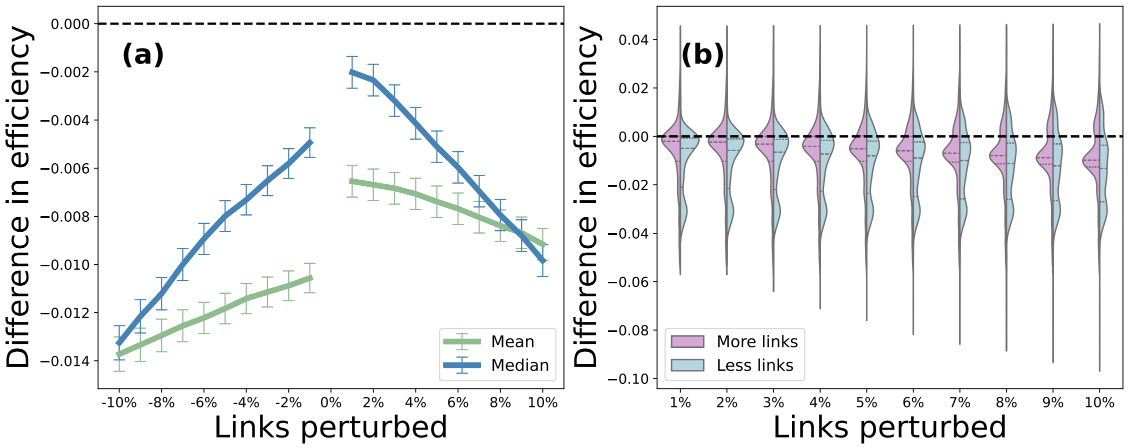


**Supplementary Figure 14: Effect of link-perturbation magnitude on network efficiency.** Change in network efficiency, $\Delta\eta$, after randomly adding or removing links from empirical ecological networks. For each network, we generated perturbed networks by modifying (1%) to (10%) of the empirical number of links and computed the difference in efficiency relative to the empirical architecture. Differences were computed across a logarithmically spaced range of propagation scales, ($\tau \in [10^{-2}, 10^2]$), and then averaged over $\tau$. Negative values indicate that the perturbed network has lower efficiency than the empirical network. **(a)** mean and median $\Delta\eta$ across networks as a function of signed perturbation magnitude, with negative values corresponding to link removals and positive values to link additions. Error bars indicate standard errors across networks. **(b)** distribution of $\Delta\eta$ values across networks for each perturbation magnitude, shown separately for link additions and removals. Across the full perturbation range, both additions and removals tend to reduce $\eta$, indicating that empirical architectures are locally consistent with high efficiency over a broad range of nearby link configurations.

#### Optimal scaling in bipartite and unipartite networks

We next asked whether the correspondence between the empirical link–species scaling and the stationary condition of the thermodynamic-efficiency approximation also holds when bipartite and unipartite networks are analysed separately. Because these two classes exhibit different empirical scaling relationships, we fitted

$$L = cS^{\gamma}$$

independently within each subset and used the corresponding value of $c$ when evaluating the finite-size stationary condition for each network.

The resulting distributions of $\gamma_{\mathrm{opt}}$ differed between the two classes in the same direction as the empirical scaling (Fig. 15). Bipartite networks yielded a mean $\langle\gamma_{\mathrm{opt}}\rangle = 1.37$, close to their empirical exponent $\gamma = 1.34$, whereas unipartite networks yielded $\langle\gamma_{\mathrm{opt}}\rangle = 1.43$, compared with an empirical exponent of $\gamma = 1.47$. Thus, the agreement between the efficiency-based stationary condition and the observed scaling is not restricted to the pooled dataset: when the empirical link–species relationship shifts between broad network representations, the corresponding efficiency-based estimate shifts accordingly.

As for the analysis of the full dataset, these values should be interpreted as finite-size consistency estimates rather than parameter-free predictions. In particular, the scaling prefactor $c$ is estimated separately from each empirical subset. Nevertheless, the ability of the approximation to reproduce the direction and magnitude of the difference between bipartite and unipartite networks provides an additional consistency check that the propagation–response-diversity trade-off is compatible with the observed variation in link–species scaling.

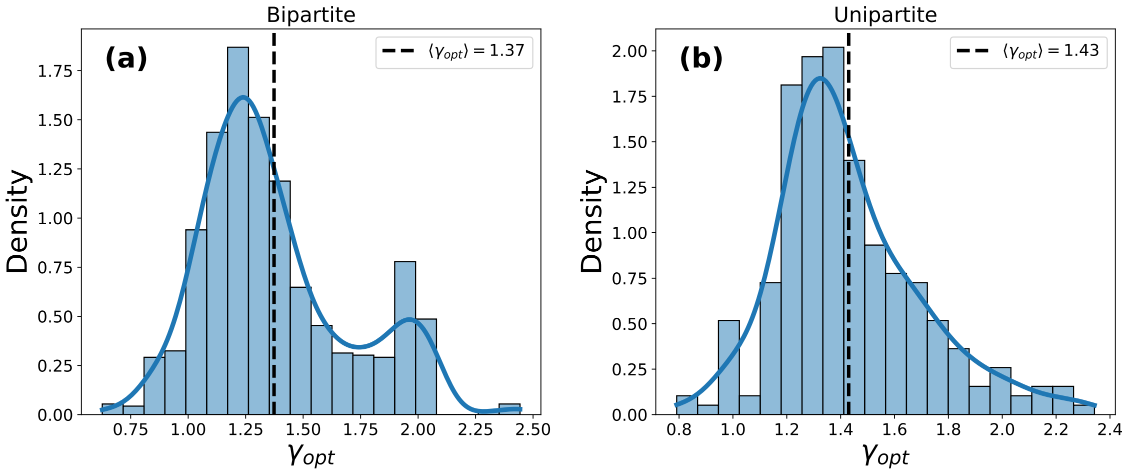


**Supplementary Figure 15: Efficiency-based scaling exponents for bipartite (a) and unipartite (b) ecological networks.** Distribution of the optimal scaling exponent, $\gamma_{\text{opt}} = \delta^* + 1$, obtained by evaluating the stationary condition of the thermodynamic-efficiency approximation separately for bipartite and unipartite networks. For each subset, the corresponding prefactor $c$ was obtained from its empirical fit $L = cS^\gamma$, while the characteristic diffusion scale of each network was taken as $\tau_{\text{diff}} = 1/\lambda_2$. The resulting mean exponents are $\langle\gamma_{\text{opt}}\rangle = 1.37$ for bipartite networks and $\langle\gamma_{\text{opt}}\rangle = 1.43$ for unipartite networks, close to the corresponding empirical estimates $\gamma = 1.34$ and $\gamma = 1.47$, respectively. The agreement shows that the thermodynamic approximation captures not only the intermediate scaling regime of the pooled dataset, but also the shift in link–species scaling between the two broad network representations.
